# Age-dependent structural reorganization of utricular ribbon synapses

**DOI:** 10.1101/2022.12.19.521049

**Authors:** Susann Michanski, Timo Henneck, Mohona Mukhopadhyay, Anna M. Steyer, Paola Agüi Gonzalez, Katharina Grewe, Peter Ilgen, Mehmet Gültas, Eugenio F. Fornasiero, Stefan Jakobs, Wiebke Möbius, Christian Vogl, Tina Pangršič, Silvio O. Rizzoli, Carolin Wichmann

**Affiliations:** Molecular Architecture of Synapses Group, Institute for Auditory Neuroscience, InnerEarLab and Center for Biostructural Imaging of Neurodegeneration, University Medical Center Göttingen, D-37075 Göttingen, Germany; Collaborative Research Center 889, University of Göttingen, D-37075 Göttingen, Germany; Multiscale Bioimaging Cluster of Excellence (MBExC), University of Göttingen, D-37075 Göttingen, Germany; Biology Bachelor Program, University of Göttingen, D-37075 Göttingen, Germany; Experimental Otology Group, InnerEarLab, Department of Otolaryngology and Institute for Auditory Neuroscience, University Medical Center Göttingen, D-37075 Göttingen, Germany; Electron Microscopy-City Campus, Department of Neurogenetics, Max Planck Institute for Multidisciplinary Sciences, D-37075 Göttingen, Germany; Center Nanoscale Microscopy and Molecular Physiology of the Brain (CNMPB), University of Göttingen, D-37075 Göttingen, Germany; Department for Neuro- and Sensory Physiology, University Medical Center Göttingen, D-37075 Göttingen, Germany and Center for Biostructural Imaging of Neurodegeneration, BIN, D-37075 Göttingen, Germany; Clinic of Neurology, University Medical Center Göttingen, D-37075 Göttingen, Germany; Department of NanoBiophotonics, Max Planck Institute for Multidisciplinary Sciences, D-37077 Göttingen, Germany; Fraunhofer Institute for Translational Medicine and Pharmacology ITMP, Translational Neuroinflammation and Automated Microscopy TNM, 37075 Göttingen, Germany; Faculty of Agriculture, South Westphalia University of Applied Sciences, D-59494 Soest, Germany; Presynaptogenesis and Intracellular Transport in Hair Cells Group, Institute for Auditory Neuroscience and InnerEarLab, University Medical Center Göttingen, D-37075 Göttingen, Germany; Auditory Neuroscience Group, Institute of Physiology, Medical University Innsbruck, A-6020 Innsbruck, Austria

**Keywords:** Aging, ribbon synapse, synaptogenesis, utricle, vestibular hair cells

## Abstract

In mammals, spatial orientation is synaptically-encoded by sensory hair cells of the vestibular labyrinth. Vestibular hair cells (VHCs) harbor synaptic ribbons at their presynaptic active zones (AZs), which play a critical role in molecular scaffolding and facilitate synaptic release and vesicular replenishment. With advancing age, the prevalence of vestibular deficits increases; yet, a direct link to the functional decline of VHC ribbon synapses remains to be demonstrated. To address this issue, we investigated the effects of aging on the ultrastructure of the ribbon-type AZs in murine utricles using various electron microscopic techniques and combined them with confocal and super-resolution light microscopy as well as metabolic imaging up to one year of age. In older animals, we detected predominantly in type I VHCs the formation of floating ribbon clusters. Our findings suggest that VHC ribbon-type AZs undergo dramatic structural alterations upon aging.

## Introduction

Sensory receptor cells in the eye and inner ear are equipped with specialized presynaptic organelles – synaptic ribbons – that tether dozens to hundreds of synaptic vesicles (SVs). This system enables indefatigable quantal release of the neurotransmitter glutamate (Matthews and Fuchs, 2010; Moser et al., 2006; Moser et al., 2019; Rutherford and Moser, 2016; Sterling and Matthews, 2005).

The mammalian vestibular system contains multiple sensory organs, the utricle and the saccule for sensing of linear acceleration and gravity, and the ampullae in the semicircular canals for the detection of head rotation (Li et al., 2016). Each of the vestibular organs harbors a sensory epithelium carrying two types of vestibular hair cells (VHCs) (Wersall, 1956). Ribbons decorate the active zones (AZs) of both, calyceal synapses of type I as well as conventional bouton synapses of type II VHCs. Morphologically, vestibular ribbons are highly variable in size, shape and number depending on the species, hair cell type and location (Eatock and Lysakowski, 2006). However, in comparison to auditory and visual ribbon synapses, relatively little is known about the synaptic architecture of vestibular ribbon synapses. Since the decline of vestibular function starts already at the age of 40 (Agrawal et al., 2009; Bermúdez Rey et al., 2016) and vestibular deficits can significantly impair the quality of life of affected patients, detailed longitudinal analyses of VHC ultrastructure upon maturation and aging are essential, but to date surprisingly scarce.

As evident by decreased amplitudes alongside increased latencies and thresholds of ocular vestibular-evoked myogenic potentials (oVEMPs) in humans (Agrawal et al., 2012; Iwasaki et al., 2008; Piker et al., 2011; Rosengren et al., 2011), vestibular function declines with age. This is in line with the observation of increased occurrences of vertigo, recurrent dizziness, nausea, disorientation, blurred vision, imbalance, and increased risk of falling in the elderly (Agrawal et al., 2012; Neuhauser, 2007; Neuhauser et al., 2005). In this context, it remains to be clarified, which cells or subcellular structures undergo age-related deterioration in the utricular system as well as other vestibular organs.

As suggested by several studies in rodent cochlear hair cells, ribbon synapse abundance, size and shape are connected to functional changes (Michanski et al., 2019; Peineau et al., 2021; Sobkowicz et al., 1982; Sobkowicz et al., 1986; Stamataki et al., 2006; Wong et al., 2014), as well as the morphology of AZ-proximal mitochondria (Stamataki et al., 2006). Furthermore, ribbon synapse dysfunction or degeneration appear to present major factors underlying age-related hearing loss (Huang et al., 2012; Peineau et al., 2021; Stamataki et al., 2006; Wong et al., 2014). Whether a similar correlation exists in VHCs is largely unknown, but it is tempting to speculate that disruption of the presynaptic energy supply, for example during aging, may negatively influence synaptic performance. Mitochondria are important for energy supply and Ca^2+^ buffering at synapses and thus play a prominent role in neuronal homeostasis (Devine and Kittler, 2018). Moreover, recent work in zebrafish lateral line neuromast hair cells indicated an important role of synaptic mitochondria in ribbon size regulation during development and synaptic function as well as structural integrity upon maturation (Wong et al., 2019).

Therefore, we set out to analyze the impact of maturation and aging on the murine utricular VHC synapses by quantifying various aspects of presynaptic AZ structure at ages ranging from 1 week to 11 months and beyond. To do so, we combined transmission electron microscopy (TEM) with nanoscale secondary ion mass spectrometry (NanoSIMS), electron tomography, focused ion beam-scanning electron microscopy (FIB-SEM) as well as confocal and stimulated emission depletion (STED) microscopy.

We found that, while AZs of both types of VHCs contained ribbons of various shapes and sizes, type I VHCs were dominated by elongated ribbons upon maturation. Moreover, type I VHCs progressively formed complex cytosolic ribbon clusters that often were void of any physical contact to the AZ membrane along with enlarged mitochondria. In contrast to previous reports describing such ribbon clusters in VHCs (Favre et al., 1986; Park et al., 1987; Sans and Scarfone, 1996; Woods and Park, 1987), we now combined a detailed presynaptic morphometric analysis with isotopic labeling to additionally establish ribbon turnover rates in elderly VHCs. This analysis revealed that floating ribbons seem to be newly-synthesized in comparison to attached ribbons, while both ribbon populations appeared much ‘younger’ than other VHC organelles such as mitochondria. In contrast, type II VHCs showed a relatively stable AZ architecture with comparable ribbon counts and shapes, as well as mitochondrial sizes independent of aging.

In summary, our data suggests that the presynaptic architecture of VHCs – in particular in type I VHCs – retains its ability to undergo active morphological changes in adult stages. Our findings support the hypothesis that new VHC ribbons are continuously formed, but fail to attach at the presynaptic AZ and hence accumulate in its proximity. Forming clusters, they could serve as a reserve pool of ribbons when needed, or alternatively, may represent pathological accumulation of ribbons rendered incapable of attaching at the presynaptic membrane.

## Results

### Upon maturation, the number of membrane-attached synaptic ribbons per presynaptic AZ decreases

To investigate maturation and aging of utricular VHC ribbon synapses, we performed a detailed ultrastructural analysis of type I and type II VHCs in mice of five age groups (postnatal day (P)9, P15, P20, 3 months, and 9-11 months). This covered the developmental period of synaptogenesis, through maturation and ultimately to the beginning of a functional decline. Behavioral studies of balance reflexes in rats revealed that the vestibular system is functionally mature around postnatal day (P)16 (Hård and Larsson, 1975; Lannou et al., 1979). However, data from detailed physiological and morphological examinations of rodents suggest that VHC synaptic connectivity does not reach a fully mature state until 3-4 weeks of age (Curthoys, 1979; Dechesne et al., 1986; Jamon, 2014; Kawamura et al., 1998). In order to distinguish between type I and II VHCs in immature P9 animals (with not yet fully developed calyces), only VHCs with clearly identifiable surrounding calyx or afferent bouton contacts were considered for analysis.

We first analyzed AZs of conventionally-embedded utricles that contain ribbons with clear attachments via a presynaptic density (PD) to the AZ membrane. In random 2D ultrathin sections, we determined the abundance, shape, and size of membrane-attached ribbons, analogous to our previous study on cochlear inner hair cells (IHCs) (Michanski et al., 2019). In immature VHCs of both types, we frequently found more than one ribbon per AZ (Fig. 1A-G), whereas, in mature VHCs, typically, a single ribbon was attached at the AZ plasma membrane (Fig. 1A, D). Additionally, we categorized synaptic ribbon shapes (Fig. 1H), since it is well established in auditory IHCs that ribbon morphology changes drastically from rather round shapes prior to hearing onset to elongated and droplet-shaped ribbons in adulthood (Michanski et al., 2019; Sobkowicz et al., 1982; Wong et al., 2014). Interestingly, upon maturation, type I VHCs displayed an apparent increase in the abundance of elongated ribbons (on the expense of droplet, wedge and round-flat ribbons). Conversely, ribbon shape distributions in type II VHCs remained with some variability much more uniform across different ages (Fig. 1H; Table S1). Our 2D TEM data further revealed remarkably stable and largely comparable mean synaptic ribbon areas in the membrane-attached ribbon populations of type I and type II VHCs as well as between the different age groups (for quantification parameters see materials and methods; Fig. S1A). Nevertheless, there was a trend of type I VHC ribbons towards enlarged heights across all age groups in comparison to type II VHC ribbons, which goes along with the observation of higher percentage of elongated ribbons in type I VHCs (Fig. 1I-K; Table S2).

**Fig. 1:**
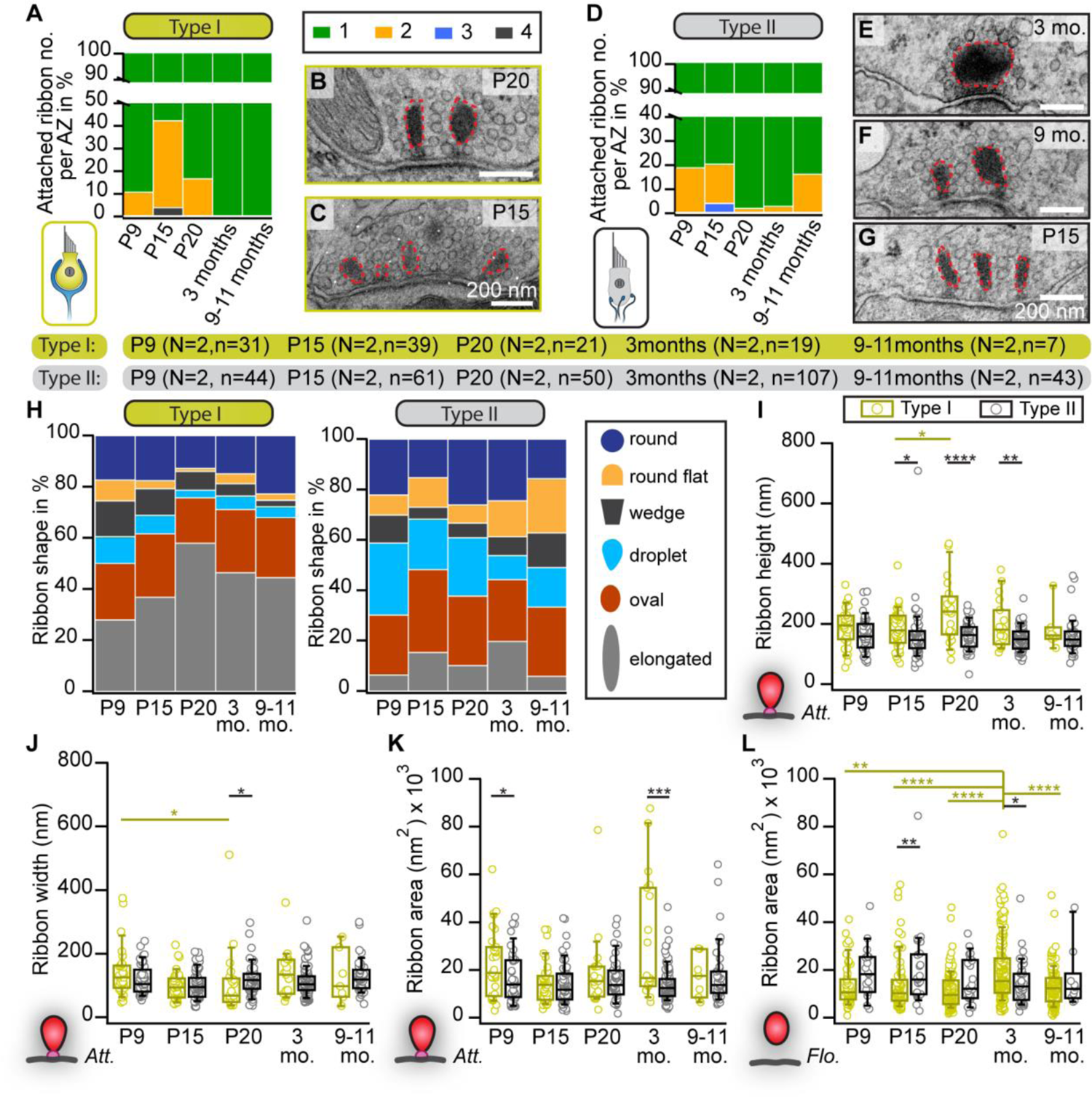
Changes in the abundance and shape of VHC ribbons upon maturation and aging. (A, D) Analysis of the number of membrane-attached synaptic ribbons reveals a larger proportion of multiple ribbons per AZ in immature type I (A) and type II (D) VHCs compared to mature ages. (B, C, E-G) Examples of electron micrographs depicting either multiple or single synaptic ribbons (highlighted with red dashed lines) in type I (B, C) and type II (E-G) VHCs. (H) Proportion of ribbon synapses categorized into different types of shapes, respectively for all quantified ages. (I-L) Box plots show random section quantifications of the ribbon size measurements for all age groups. No prominent changes regarding the ribbon area, height, and width could be observed in older animals. N = animal number, n = ribbon number. Significant differences between two groups were analyzed with a two-tailed t-test or Mann– Whitney Wilcoxon test. For multiple comparisons, ANOVA followed by the post-hoc Tukey or Kruskal-Wallis (KW) test followed by NPMC test was performed. For more detailed information see also Table S2.

To our surprise, in contrast to mature auditory IHCs, we observed several ‘floating’ synaptic ribbons – i.e., ribbons lacking any apparent contact with the plasma membrane – throughout all age groups and in both VHC types (Fig. S1B; Table S3). While no major systematic differences in floating ribbon size could be observed in either VHC type or between age groups, floating ribbons from 3-month-old type I VHCs displayed significantly larger ribbon areas in comparison to type I VHCs in other age groups (Fig. 1L; Table S2).

### Progressive increase in the number of floating ribbons and ribbon cluster formation in type I VHCs

A characteristic feature of immaturity in cochlear IHCs is the presence of round, free-floating ribbon precursors, which attach to the AZ membrane or might fuse with membrane-anchored ribbons during maturation (Michanski et al., 2019; Sobkowicz et al., 1982; Wong et al., 2014). Conversely, floating ribbons appear to persist in VHCs independent of their developmental stage (Fig. 2A, B).

**Fig. 2:**
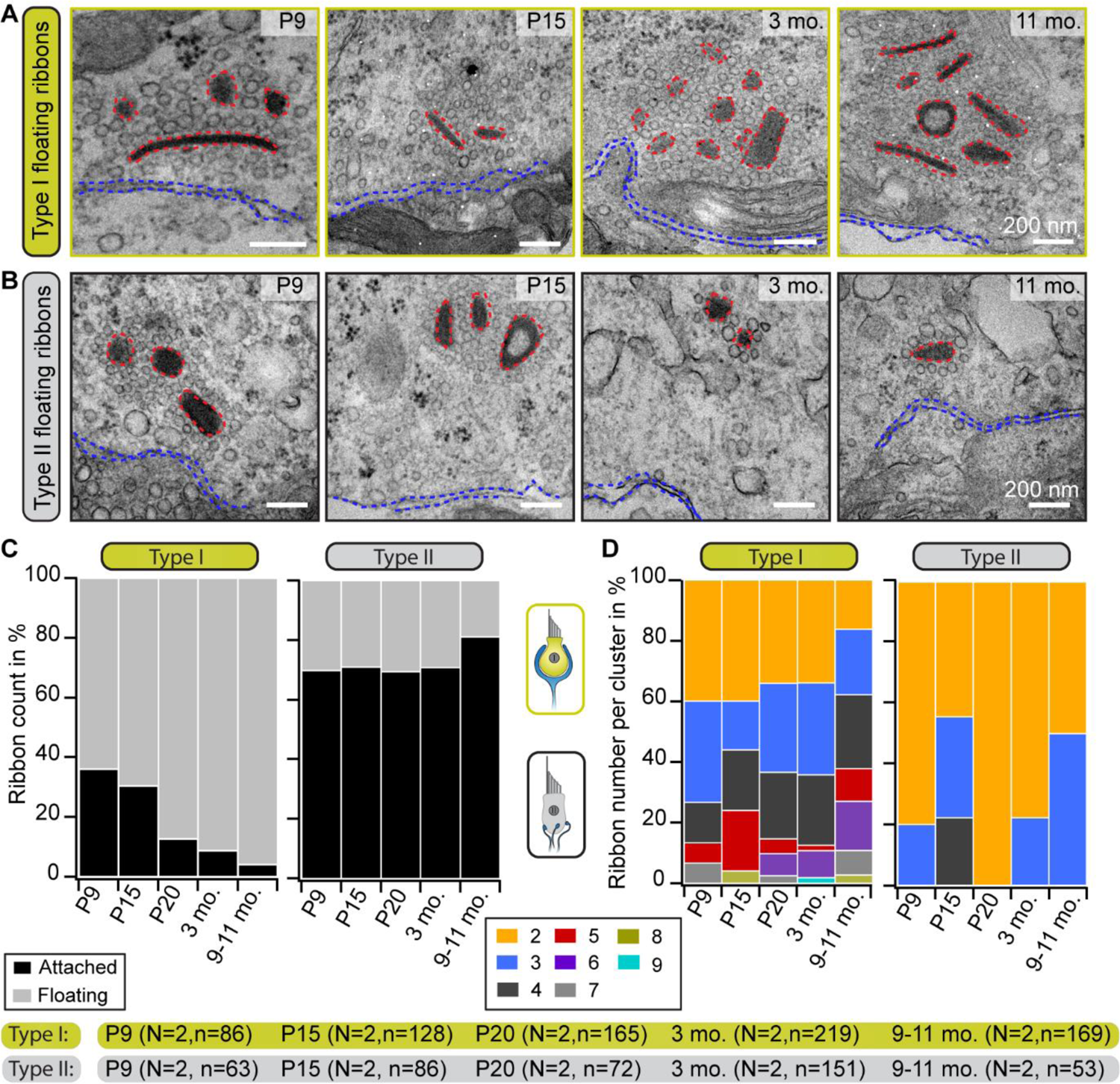
Progressive accumulation of floating ribbons in type I VHCs. (A, B) Representative electron micrographs showing single and multiple floating ribbons (red dashed lines) in type I (A) and type II (B) VHCs. All depicted ribbons appear to lack the presynaptic contact to the HC membrane (blue dashed lines). (C) Graphs representing the proportions of floating and membrane-attached ribbons during maturation and aging. Mature type I VHCs possess predominantly floating ribbons, whereas type II VHCs exhibit mainly attached ribbons throughout all investigated ages. (D) Quantification of the total number of ribbons per cluster reveals the occurrence of large ribbon clusters in both VHC types. With advancing age, the ribbon count per cluster increases in type I VHCs but slightly decreases in type II VHCs. N = animal number, n = ribbon number.

To now further characterize these organelles in greater detail in both VHC types, a range of different ultrastructural approaches was employed. In 2D random sections, we observed that, while the fraction of floating ribbons in type I VHCs dramatically increased with age – ultimately constituting ∼95% of all AZ-proximal ribbons in the 9-11 months age group – (Fig. 2C; Table S3), the proportion of floating ribbons in type II VHCs remained remarkably stable at ∼20-30%. When measuring the nearest distance between these floating ribbons and the plasma membrane, we found appositions predominantly ∼200 nm from the AZ membrane, with generally tighter spacing in type II VHCs for almost all age groups (Fig. S1D; Table S8). This finding may be indicative of a progressive age-dependent anchoring deficit – especially in type I VHCs.

These floating ribbons then further accumulated into clusters (≥ 2 ribbons) with increasing complexity and often – but not always – resided in direct proximity to membrane-attached ribbons at the AZs of both VHC types. In type I VHCs, such clusters were composed of up to nine ribbons (Fig. 2A, D; Fig. S1C), whereas a maximum of four – but mostly a pair of ribbons per cluster – were observed in type II VHCs analyzed from random sections (Fig. 2B, D; Table S3). In type I VHCs, the ribbon clusters grew continuously from P20 onwards, while a similar trend was not observed in type II VHCs. There, we even detected a decrease of cluster ribbon content at P20 followed by a slight increase at older ages (Fig. 2D). A striking increase in the proportion of single ribbons was observed in type II VHCs, whereas in type I VHCs, synaptic ribbons in clusters clearly dominated at older ages (Fig. S1E; Table S3).

Since floating and attached ribbons alike exhibited a translucent core in some cases (Fig. 2A, B and Fig. S1C), we quantified the frequency of occurrence of this phenomenon. This special ribbon morphology has previously been described in several vestibular and cochlear studies, but mostly in adult tissue (Favre and Sans, 1979; Liberman, 1980; Michanski et al., 2019; Park et al., 1987; Sjöbäck and Gulley, 1979; Sobkowicz et al., 1982; Stamataki et al., 2006). Thus, it has been proposed that this feature might indicate ‘old’ ribbons prior to their degradation. Evaluation of floating and attached VHC ribbon counts with a translucent core in our study revealed no age-dependent increase in this ribbon feature (Fig. S1F). Therefore, we hypothesize that ribbon size – not structural ‘age’ – determines the formation of a translucent core.

Since the nature of 2D-analysis may lead to a systematic under-estimation of ribbon attachment, we next verified the abundance of free-floating ribbons and their accumulation in larger clusters, by performing electron tomography. This 3D approach confirmed the occurrence of ribbon clusters consisting either of exclusively floating ribbons (Movie 1) or floating together with some membrane-attached ribbons (Movie 2), predominantly in type I VHCs vs. mainly attached single ribbons in mature type II VHCs (Movie 3).

**Movie 1-3: Electron tomography with 3D segmentation of ribbon cluster and single attached ribbons**.

Tomograms and their corresponding 3D models of mature (3 months) type I/II VHCs depicting several disc-shaped floating ribbons in type I VHCs and typical single attached ribbons in type II VHCs. The different SV pools, membrane-proximal (MP)-SVs and ribbon-associated (RA)-SVs, are defined as displayed in Fig. S1A.

Red: ribbon, magenta: presynaptic density, dark blue: postsynaptic density, light blue: HC membrane, yellow: SVs, orange: MP-SVs, green: RA-SVs, brown: SV not associated with the ribbon but part of the cluster “cloud”. Scale bars: 200 nm.

**Movie 1:** https://owncloud.gwdg.de/index.php/s/bRolkq37mZxUCy5

**Movie 2:** https://owncloud.gwdg.de/index.php/s/DfmaoBlUGUHHJiA

**Movie 3** https://owncloud.gwdg.de/index.php/s/u4rJwBpUWlzGJj3

To exclude the possibility that the preparation method of the delicate tissue could have caused putative artifacts leading to a higher frequency of floating ribbons, we applied an alternative sample preparation method. Here, slow fixative perfusion of the inner ear followed by utricle dissection yielded similar results as with our standard preparation method. Moreover, the previously reported age-related progressive genetic hearing loss in C57BL/6J mice (Henry and Chole, 1980; Hequembourg and Liberman, 2001; Ison and Allen, 2003; Johnson et al., 1997; Li and Borg, 1991; Mikaelian et al., 1974; Shnerson and Pujol, 1981; White et al., 2000; Willott, 1986) due to the Cdh23 mutation, did not strongly affect vestibular function and morphology does not seem to be altered (Mock et al., 2016). Nevertheless, to exclude the possibility of mouse background related structural changes in VHCs, we further examined utricles of adult (6-month-old) CBA/J wild-type mice. These mice maintain relatively normal hearing until they reach senescence (Shone et al., 1991; Spongr et al., 1997; Walton et al., 1995; Willott, 1986; Willott et al., 1988a; Willott et al., 1988b). Electron micrographs of random and serial sections from CBA/J mice confirmed an age-dependent increase in the frequency of floating ribbons as well as the presence of complex ribbon clusters in type I VHCs (Fig. S2).

### Molecular constituents of ribbon clusters

The extremely high percentages of cytoplasmically floating ribbons – especially in type I VHCs (∼90% already at P20; Fig. 2C) – led to the question whether floating and membrane-attached ribbons in type I and II VHCs are different in their molecular composition from attached ribbons in other HC types. For example, in cochlear IHCs, it could previously be shown that genetic loss of the large structural scaffold bassoon prevented adequate ribbon anchoring at the AZ membrane (Khimich et al., 2005). Therefore, immunohistochemical stainings of the main ribbon components RIBEYE and piccolino as well as the molecular anchor bassoon were performed in order to determine the localization of these proteins. We detected piccolino on floating ribbons as well as membrane-attached ribbons being part of a ribbon cluster in both VHC types (Fig. 3). A qualitative assessment of the images suggested the presence of clusters in both young and older animals, while some very large clusters were only observed in the older animals. Fig. 3F and L illustrate large ribbon clusters, which were only observed in 32-week-old animals. It is also noteworthy that panel F and the first image in lower panel D (Fig. 3; asterisks) depict ribbons from 32-week-old mice without a nearby bassoon spot, which could indicate that these ribbons lack a membrane-anchor and thus likely represent floating ribbons.

**Fig. 3:**
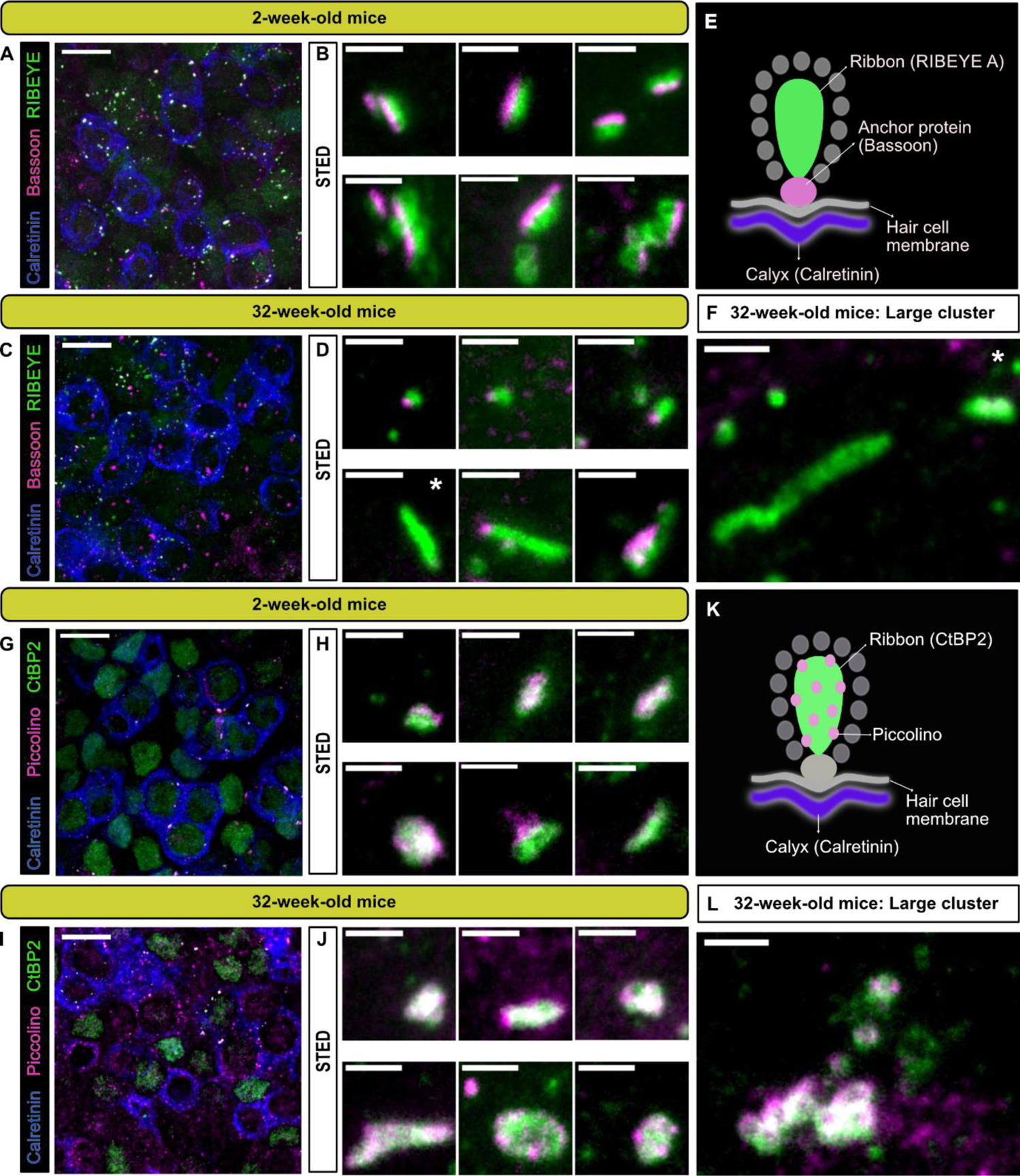
Localization of the presynaptic AZ proteins bassoon and piccolino at type I VHCs ribbon synapses. (A,C,G,I) 100x confocal images of the utricular striola, identified here by the presence of calretinin (blue) stained single and complex calyces, with combinations of various synaptic proteins. Ribbon proteins RIBEYE A (A,C) and CtBP2 (G,I) shown in green. Anchoring protein bassoon (B,D) and ribbon-associated protein piccolino (G,I) shown in magenta. Scale bars in A, C, G, and I: 10 µm. (B,D,H,J) Top panels show representative STED images of putative single ribbons or small ribbon clusters and lower panels show images of complex clusters near the type I VHC membranes, with the respective antibody combinations (scale bars for STED images: 1 µm). (E,K) Schematic representation of the antibody combinations used for each panel in young (2 weeks) and old mice (32 weeks). (F and L) Representative STED images of particularly large ribbon clusters (scale bar: 1 µm). N = 3 animals for each combination. Note that 100x confocal images represent maximal projections of only a few optical planes for better visibility. White asterisks mark the panels depicting ribbons that seemingly lack bassoon (even in the optical planes that were omitted from projections).

The utricle is a remarkably dense tissue, with multiple layers of tightly-packed type I and type II VHCs. Especially for EM sectioning, this poses an intrinsic problem, as distinction between striolar and extrastriolar VHCs – even of the same VHC type – can be immensely difficult, but may influence our results and interpretation. Hence, we decided to perform a basic comparative analysis of ribbon parameters across aging type I VHCs from striolar and extrastriolar regions (Fig. S3). For this purpose, we bred *Ai14-Neurog1-creER^T2^* knock-in mice (Druckenbrod and Goodrich, 2015; Koundakjian et al., 2007; Madisen et al., 2010) to sparsely express the red fluorescent protein tdTomato in a small subset of vestibular ganglion neurons. This strategy enabled us to trace individual neurons and clearly identify their afferent fibers with VHCs. By combining this with immunohistochemistry, we could then – apart from cellular location in the fixed utricle – either select individual VHCs with calretinin-positive and tdTomato-positive calyces (as indicative for calyx-only fibers in the striola) or calretinin-negative, but tdTomato-positive calyces (representative of the calyceal endings of dimorphic fibers in the extra-striolar region) and subject them to quantitative analysis. Using this approach, we could unambiguously and accurately establish ribbon counts and volumes of individual type I VHCs that were randomly labeled in either the striolar or extrastriolar regions and extended this analysis for three cohorts covering a time period from 2-3 weeks to 1.5 years of age. In line with previous work (Wan et al., 2019) and independent of location, we observed a decline in overall ribbon spots (Fig. S3B, D). Yet, while we found a statistically significant increase in ribbon spot size with advancing age in the striolar region, such effects could not be detected in the extrastriolar region in these experiments (Fig. S3B’, D’).

While this approach offers interesting new insights, it also comes with a range of drawbacks: for example, one has to take into account that due to the limited lateral and axial resolution of confocal microscopy (the latter also applies to 2D-STED), small and dense ribbon clusters – as observed in our 2D TEM data – will appear as one large ribbon in immunofluorescence experiments. Therefore, the true ribbon number per VHC is likely to be underestimated. Even in our STED analysis, large appearing ribbons or clusters might still not be completely resolved as separated entities, and this way might escape correct quantification under confocal microscope. Moreover, due to the very close spacing between floating ribbons and the AZ plasma membrane – predominantly below the diffraction limit of ∼200 nm – parameters such as synaptic engagement could not reliably be investigated using this method.

Taken together, while all employed 2D techniques revealed a range of novel insights into VHC synaptic morphology and age-dependent structural alterations at VHC AZs, it became clear that a large-scale high-resolution 3D method is required to determine the number of ribbons and ribbon clusters per cell and their true dimensions in 3D.

### 3D volume analysis of the basolateral compartment of VHCs reveals larger ribbons in mature type I VHCs

In order to investigate ribbon clusters in more contextual detail, we performed focused ion beam – scanning electron microscopy (FIB-SEM) to obtain 3D information of the entire basolateral compartment of individual VHCs. These data could then be used to assess various ultrastructural parameters, including presynaptic ribbons alongside individual SVs and the postsynaptic innervation – at lateral resolutions of 3 or 5 nm and an axial resolution of 5 nm. Thereby, morphological parameters including ribbon number, size and the location of ribbon clusters can be visualized (Fig. 4). Since FIB-SEM is a very complex, expensive and time-consuming method, we focused only on two representative ages: P15 for a maturing age group, and 8-month-old for an adult stage.

**Fig. 4:**
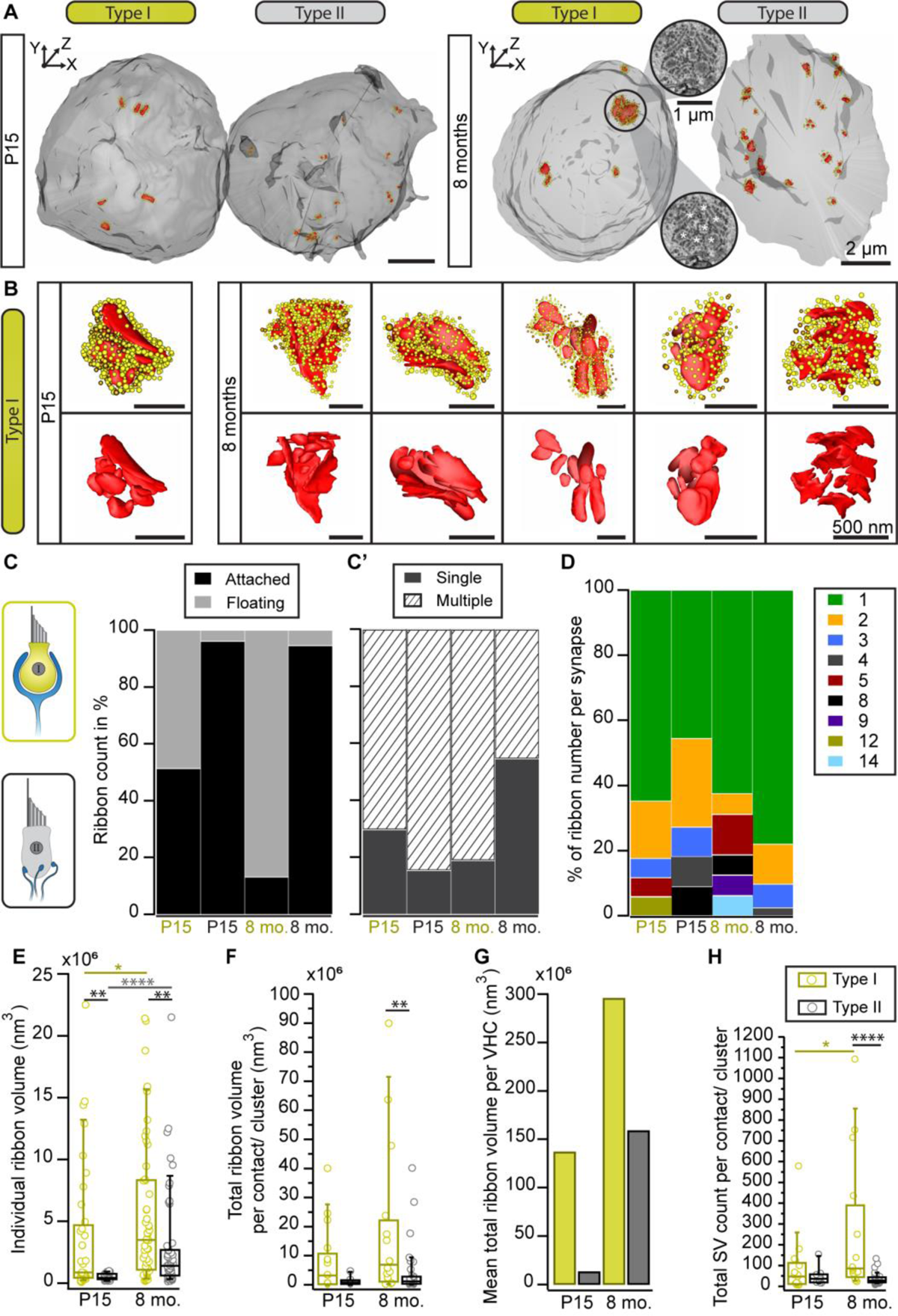
Ribbon cluster abundance in large volume data. (A) 3D models of type I and II VHCs from P15 and 8-month-old mice using FIB-SEM (top view). The P15 data set derived from type I VHCs with a complex calyx. The VHC bodies (light gray) contain numerous ribbon synapses (red) tethering SVs (yellow: ribbon-associated, orange: peripheral SVs). (B) Individual ribbon cluster segmentations of immature and mature type I VHCs. Upper row shows ribbon clusters with SVs, lower row displays the respective segmentation without SVs. (C, C’) Bar graphs showing the distribution of attached vs. floating and single vs. multiple ribbons per VHC type and age group. (D) Analysis of the total ribbon count per cluster reveals more ribbon numbers in type I VHCs. (E-G) Ribbon size measurement data from both VHC types and age groups (Student’s t-test or Mann-Whitney-Wilcoxon test). (H) Boxplot illustrating the quantification of SV numbers per contact or cluster (*\*P* = 0.0178, \*\*\*\**P*<0.0001 Mann-Whitney-Wilcoxon test). Note, the high SV count in type I VHCs with >1000 SVs associated with a single cluster. For more detailed information see also Table S6.

Consistent with the 2D random section analyses and electron tomography data, 3D segmentations from FIB-SEM volumes revealed the occurrence of ribbon clusters with increased frequencies in type I VHCs (Fig. 4A, B; Movie 4 and 5) – particularly in mature animals. In fact, the clusters were larger than expected from the 2D TEM data or the immunohistochemistry and most ribbons within a cluster were indeed floating in the 8-months age group (Fig. 4C; Table S4). Conversely, type II VHCs exhibited predominantly attached ribbons in both age groups, which might be a result of a clear assignment – floating vs. attached – which is only possible in large 3D reconstructions (Fig. 4C; Table S4; Movie 6 and 7). Moreover, both VHC types showed higher numbers of multi-ribbon (attached and floating together) AZs per synaptic contact site in older ages. The AZs typically contain one to two (in type II VHCs) attached ribbons, which, in particular in old type I VHCs, are often surrounded by a cluster of floating ribbons. As mentioned above, in comparison to our 2D analysis, the ribbon counts per cluster were larger at both investigated ages (Fig. 4C’; Table S4) – with a maximum of 14 ribbons per cluster observed in a type I VHC (Fig. 4D; Table S4). While type II VHCs had on average more ribbon clusters per cell (P15 = 9 clusters, n = 1 VHC; 8 months = 5 clusters, n = 2 VHCs), fewer but more complex ribbon clusters were found in type I VHCs (P15 = 2 clusters, n = 3 VHCs; 8 months = 1.5 clusters per VHC; n = 4 VHCs). The average number of ribbons per cluster rose from P15 with 4.8 to 7.2 at 8 months in type I VHCs. In contrast, the average ribbon number per cluster remained constant in type II VHCs with advancing age (P15 = 2.4 ribbons, 8 months = 2.5 ribbons), largely consistent with our 2D data (Fig. 2). In addition, there are fewer ribbons per cell observed in type I VHCs at both age groups (Table S5) with fewer single membrane-attached ribbons (on average 2 ribbons) in 8-month-old type I VHCs compared to 8-month-old type II VHCs (on average 15 ribbons). For more detailed information about ribbon counts per individual VHC please refer to Table S5, which provides absolute ribbon counts for the different ribbon categories.

Quantifications of ribbon volumes revealed an age-dependent increase in individual ribbon volumes as well as the total ribbon volume per VHC (Fig. 4E-G; Table S6). Thus, the FIB-SEM data provide new insights into the differences between type I and II VHCs regarding the dimensions and counts of clusters as well as ribbon volumes. The 2D random section data showed only a slight trend towards larger ribbons in type I VHCs (Fig. 1I), while large volume data enabled accurate quantification of individual ribbon volumes (Fig. 4E-G; Table S6) alongside the number of cluster-associated SVs. In fact, a single cluster could accumulate >1000 SVs in our data sets (Fig. 4H). Moreover, the summed ribbon volume per VHC indicated an age-dependent increase in ribbon material in both VHC types, with type I VHCs containing strikingly more ribbon material than type II VHCs in both age groups (Fig. 4G).

**Movie 4 and 5: FIB-SEM visualizations of type I VHCs with 3D segmentation of single ribbons and ribbon clusters.**

Movies scanning through the representative FIB-SEM z-stacks of P15 (Movie 4) and 8-month-old (Movie 5) type I VHCs, showing 3D reconstruction of the VHC contours (transparent gray), part of the nuclei (dark gray), synaptic ribbons (red) and their corresponding SVs (yellow). Movie 4 displays two neighboring P15 type I VHCs likely from the striola that are enclosed by one complex calyx.

**Movie 4:** https://owncloud.gwdg.de/index.php/s/p7cMu7Vn56Eo8IJ

**Movie 5:** https://owncloud.gwdg.de/index.php/s/3FBCClQn8qRYcpo

**Movie 6 and 7: FIB-SEM visualizations of the basolateral compartment of type II VHCs with corresponding 3D segmentations.**

Movies scanning through the FIB-SEM z-stacks of P15 (Movie 6) and 8-month-old (Movie 7) type II VHCs. The corresponding 3D models depict VHC contours (transparent gray), part of the nuclei (dark gray), innervating nerve fibers (blue), mainly single attached ribbon synapses (red) and their corresponding SVs (yellow).

**Movie 6:** https://owncloud.gwdg.de/index.php/s/7oOyQvonWaDtOc5

**Movie 7:** https://owncloud.gwdg.de/index.php/s/6IlGWtXMvdHVri5

### Ribbon clusters may act as SV reservoirs

Our observation that ribbon clusters bind a large reservoir of SVs raised the question, if these SVs can actually still contribute to the SV cycle. The large number of SVs associated with the ribbon clusters – especially in type I VHCs – might either indicate (i) on-demand recruitable SV reservoirs or (ii) SV buffers that hold SVs that were removed from the release cycle (Fig. 4B, H). To examine SVs in more detail, we first analyzed the total amount of SVs within 80 nm distance from the ribbons in our high resolution 2D TEM data set. For attached ribbons, we additionally distinguished between SVs in membrane proximity (MP-SVs: ≤ 25 nm distance from the AZ membrane and ≤ 80 nm from the presynaptic density) and ribbon-associated vesicles (RA-SVs: first row of SVs around the ribbon within 80 nm) (Fig. S4), which represent two well-studied morphological SV pools at auditory ribbon synapses and might give some indications about SV availability (Chakrabarti et al., 2018; Jean et al., 2018; Lenzi et al., 1999; Lenzi et al., 2002; Strenzke et al., 2016). Due to the missing attachment point of floating ribbons to the membrane, the ribbon orientation could not be identified and hence, only the total SV pool was analyzed for floating ribbons.

Results from floating and attached ribbons revealed a tendency towards higher numbers of total SV, MP- and RA-SV counts in type I VHCs especially at older ages, similar to the data from FIB-SEM (Fig. 5A, D; Fig. S4A, B; Table S7). However, the SV density of attached and floating ribbons, normalized to the ribbon surface area (Fig. 5B, E), as well as the fraction of MP-SV and RA-SV pools revealed no age-related differences for both VHC types (Fig. S4C, D; Table S7). Finally, SV diameters were measured since our previous study revealed a decrease in SV diameters upon development of IHC ribbon synapses (Michanski et al., 2019). Interestingly, SV diameters from attached and floating ribbons showed significantly larger SVs in type II VHCs compared to type I VHCs with no age-dependent effects (Fig. 5C, F; Table S8).

**Fig. 5:**
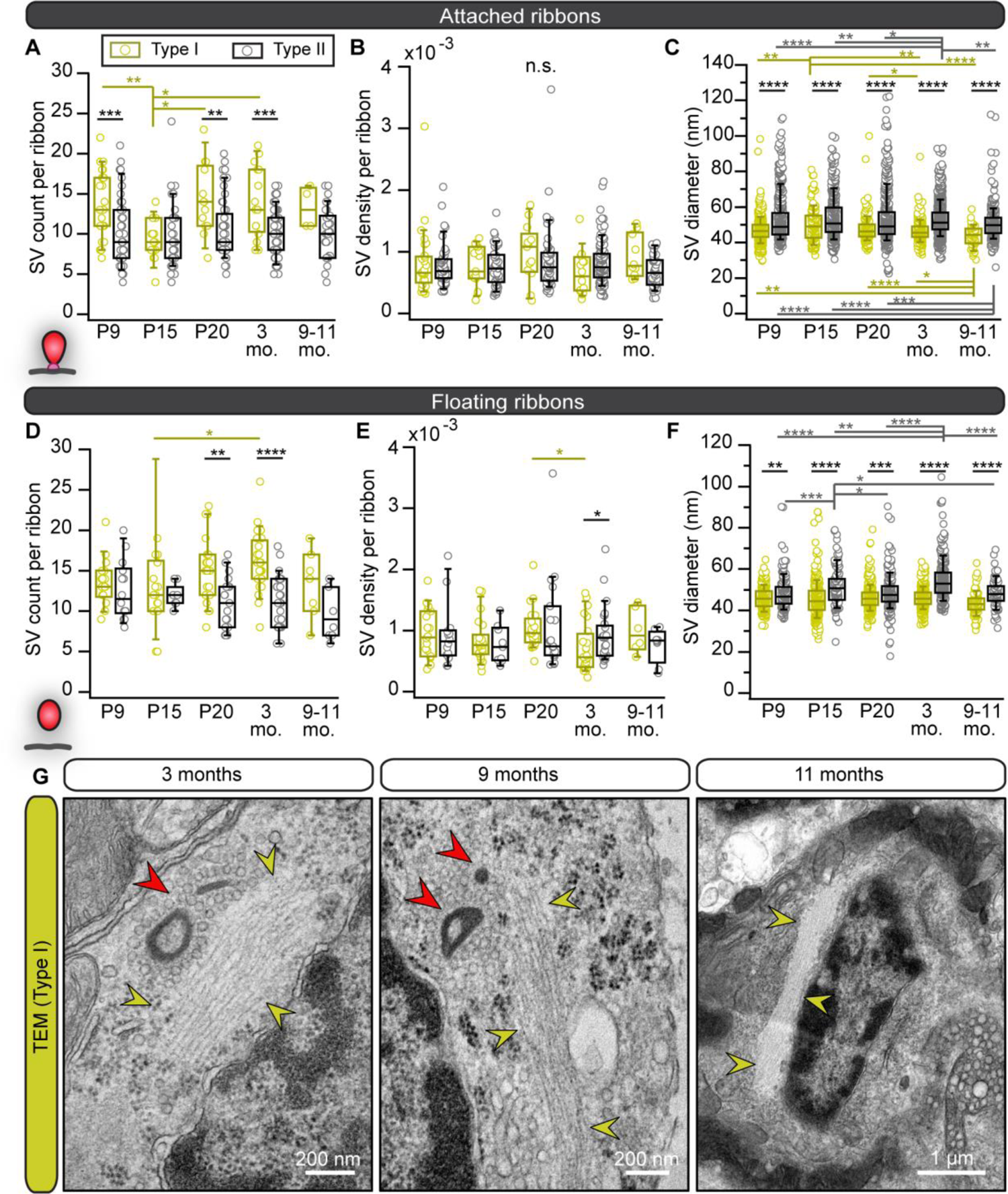
Smaller SV diameters but larger SV clouds around type I VHC ribbons close to filamentous structures. (A-F) Boxplots present the data for SV counts, their density and diameter measurements for attached and floating ribbons, respectively. (G) Electron micrographs from 2D TEM data display accumulations of filamentous cytoskeletal structures (yellow arrowheads) in type I VHCs of older ages, which were regularly detected nearby ribbon clusters (red arrowheads) with their corresponding excessive clouds of SVs. For more detailed information see Table S7 and S8.

### An extensive apico-basal microfilament network may stabilize aged type I VHCs

Lastly, from ∼3 months on, type I VHCs exhibited prominent strands of longitudinal filaments that could represent cytoskeletal elements such as microtubules or actin filaments (Fig. 5G). These strands were partially wrapped around the nucleus in a spiral manner, as segmented in our FIB-SEM data sets (Movie 8, Fig. S4F). About the function of these large cytoskeletal strands, likewise actin, can only be speculated; however, similar structures – previously termed ‘microfilaments’ – have been described in Heywood et al. (Heywood et al., 1975). If these microfilaments support ribbon (cluster) transport to the AZ remains elusive to date. Yet, in our previous study in cochlear IHCs, we described a potential involvement of microtubules in ribbon transport to the AZ due to close spatial proximity and co-localization of floating ribbon precursors with the microtubule-based motor protein KIF1a (Michanski et al., 2019). In addition, such microfilaments are likely also involved in maintaining cell shape and longitudinal stability.

**Movie 8: Filamentous network at the nucleus level can be traced in a partial FIB-SEM z-stack of an 8-month-old type I VHC.**

Movie displays a large microfilament network (highlighted with arrow heads) proximal to a ribbon cluster (highlighted by a red circle).

**Movie 8:** https://owncloud.gwdg.de/index.php/s/W3ZsvG5NWAqbCuM

### Mature type I VHCs exhibit larger mitochondria that reside closer to the AZ

Ribbons are designed for release sites with large membrane turnover and commonly operate in sensory systems characterized by tonic – probably high-rate – SV release. Continuous synaptic SV cycling upon tonic transmission might require high amounts of metabolic energy, i.e., ATP. Since production of ATP requires adequate mitochondrial function, we next assessed mitochondria abundance and morphology in our 3D FIB-SEM datasets. Indeed, in 8-month-old type I VHCs, the overall mitochondrial volume was increased four-fold as compared to the young controls with some mitochondria reaching sizes up to 100-fold larger than the average mitochondrion in the young type I VHC (volume of 4.29 x 10^7^ nm^3^; Fig. 6A-E). In contrast, much smaller differences were detected in young vs. aged type II VHCs (Movie 9 and 10). The presence of enlarged mitochondria in type I VHCs could further be observed with TEM already at 3 months of age (Fig. S5).

**Fig. 6:**
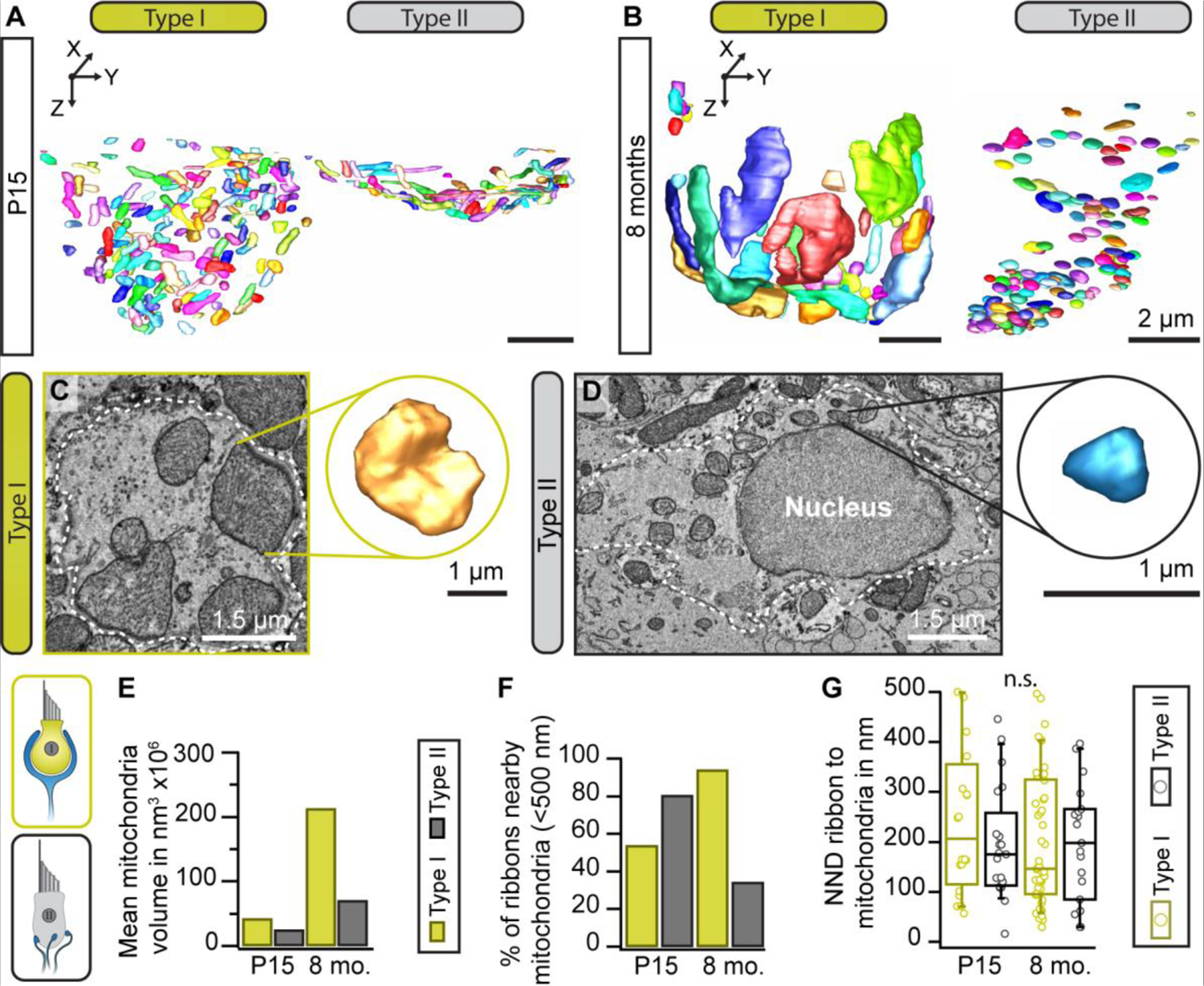
Enlargement of mitochondrial volume in type I VHCs with advancing age. (A, B) Representative 3D segmentations of mitochondria in type I and II VHCs at indicated ages. (C, D) Single sections from the large FIB-SEM datasets illustrating the mitochondrial size difference between type I and type II VHCs. Insets illustrate 3D reconstruction of the individual mitochondrion. (E) Analysis of the mean mitochondria volume presents notably larger mitochondria in mature type I VHCs. (F) Percentage of mitochondria within <500 nm of the closest ribbon in both age groups and VHC types. (G) Nearest neighbor distance (NND) of mitochondria-proximal ribbons (located <500 nm to the closest mitochondrion), *n.s. P*>0.05 (t-test).

Since mitochondria were suggested to importantly influence presynaptic activity through ATP, Ca^2+^ supply and buffering (Esterberg et al., 2016; Pickett et al., 2018; Wong et al., 2019), their number and distance to ribbons might change during maturation of synaptic machinery and structure. Indeed, our FIB-SEM data suggest a developmental rise of mitochondrial counts nearby synaptically-engaged ribbons in type I VHCs with a slight trend towards a closer proximity that was not observed in type II VHCs (Fig. 6F, G). Here, the opposite was found with fewer mitochondria nearby ribbons in older animals.

**Movie 9 and 10: 3D visualization of mitochondria from FIB-SEM z-stacks of P15 and 8-month-old type I and type II VHCs.**

Movies show 3D reconstructions of VHC contours (transparent gray), part of the nuclei (dark gray) and mitochondria (randomly colored).

**Movie 9** (P15): https://owncloud.gwdg.de/index.php/s/ICNrhNwO27HTgzm

**Movie 10** (8 months): https://owncloud.gwdg.de/index.php/s/ZyYt62UINSeGGHv

### VHC ribbons show a high turnover

Since ribbon material seems to accumulate specifically in older type I VHCs, the question remains, where the observed floating ribbons originate from. Here, two scenarios can be envisioned: (i) at the end of their lifetime, ‘old’ ribbons detach (with SVs) for adequate degradation and resorption; however, this process might get corrupted with advancing age resulting in accumulation of ribbons close to the AZ, or (i) new ribbons are constantly being formed in the cytosol and transported to the AZs; however, once there, the membrane-anchoring at the AZ might fail or become inefficient, thus leading to cluster formation. While given parts of our current data may support either scenario, we next set out to clarify this issue directly via metabolic labeling.

For this purpose, we made use of nanoscale secondary ion mass spectrometry (NanoSIMS). In NanoSIMS, a primary Cs^+^ beam irradiates the sample and causes the sputtering of secondary particles from the sample surface. These particles are partly ionized and subsequently identified by mass spectrometry. In these experiments, mice were fed for 14 or 21 days with a modified diet, in which every new nitrogen atom is provided as the heavy ^15^N stable isotope variant that, in concentrations higher than ∼0.4% can be unambiguously distinguished from the far more abundant (>99%) naturally-occurring ^14^N. This approach provides unbiased insights into the composition of the large majority of cellular structures, since not only proteins, but also nucleic acids, lipids (e.g., phospholipids as phosphatidyl-choline, -ethanol amine or –serine, and sphingolipids) and sugars (e.g., N-acetylated glycans) contain nitrogen atoms. Hence, it serves as an excellent approach to identify “old” vs. “new” material in VHCs by calculating the ^15^N/^14^N ratio. Typically, this approach allows for lateral resolutions ∼50-100 nm and axial resolutions of ∼5-10 nm. To now relate turnover rate to cellular ultrastructure, we combined NanoSIMS with 110 nm ultrathin sections of conventionally-embedded samples (Fig. 7).

**Fig. 7:**
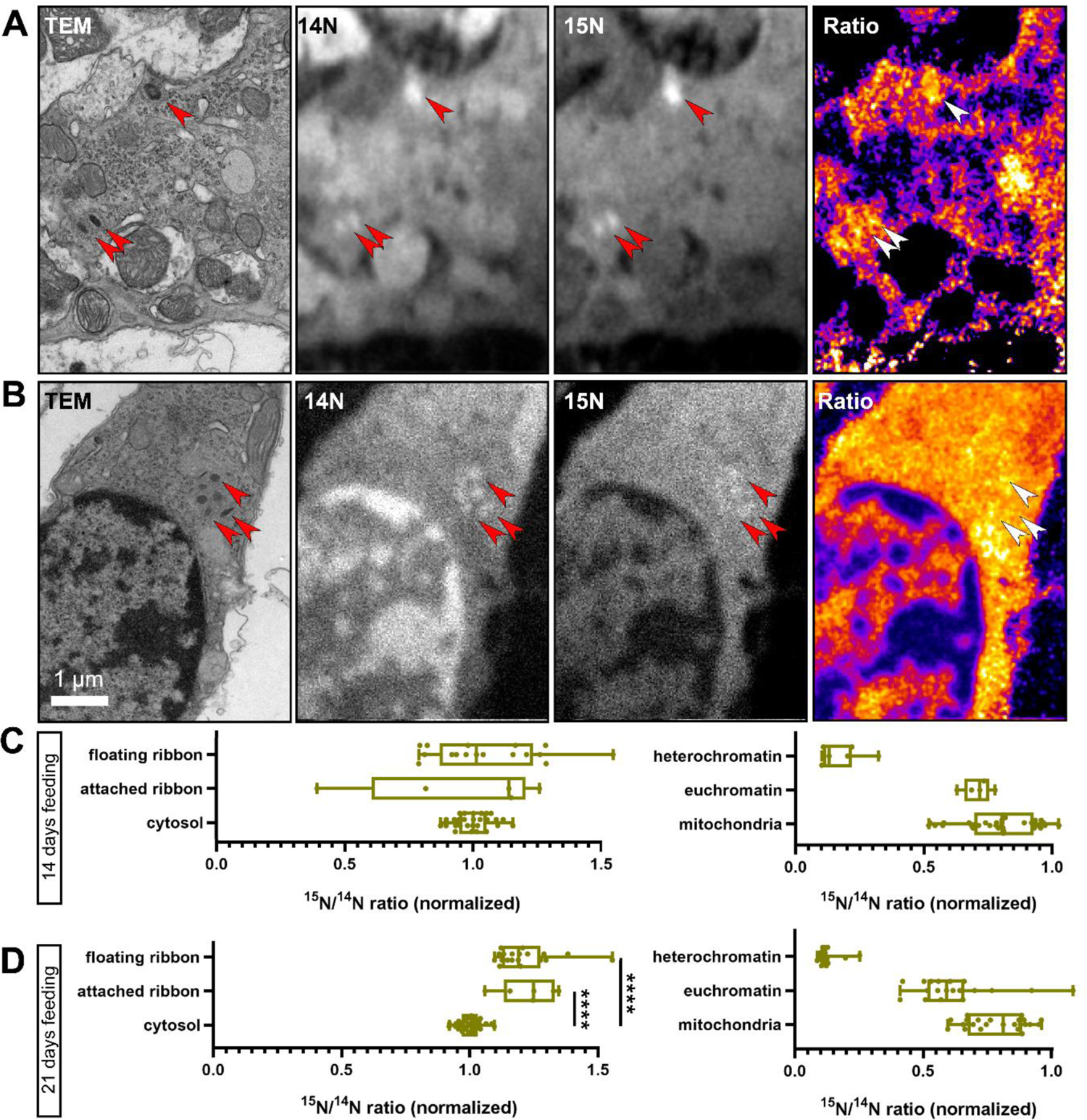
NanoSIMS analysis of VHC ribbons. (A-B) To measure protein turnover in VHCs, newly synthesized cellular components were labeled with ^15^N via a modified diet. 10-14 months old wild-type mice were subjected to this diet for 14 days, before dissecting the VHCs and imaging them first in TEM and then in NanoSIMS. The arrowheads point to several ribbons, with (A) indicating a type II VHC and (B) a type I VHC. In (A) the topmost ribbon is membrane-attached, while the other two are floating. In (B), all six ribbons are floating in the cytosol. The ^14^N and ^15^N images indicate that the protein-dense ribbons are easily visualized in NanoSIMS, as bright (nitrogen-intense) spots. The ratio images show the ^15^N (newly synthesized) signal divided by the ^14^N (old) signal. The ribbons in (A) are considerably richer in ^15^N than the ones in (B). (C) The analysis indicates the ^15^N/^14^N ratios for 13 attached ribbon and 40 floating ribbon measurements, from 15 series of NanoSIMS measurements, from 4 organs. The floating ribbons are substantially more enriched in ^15^N than mitochondria or heterochromatin (*P*<0.0001, KW test). The attached ribbons are also significantly more enriched in ^15^N than heterochromatin (*P*<0.0001, KW test). (D) A similar analysis for organs from animals fed with ^15^N for 21 days. The ribbons are now substantially richer in ^15^N than all other cellular components (*P*<0.0001; 6 to 25 measurements). The attached ribbons low in ^15^N (old) have now disappeared, suggesting that ribbons are replaced on a 1-3 week timeline. N14 days = 2 animals; N21 days = 2 animals.

Surprisingly, we found that both the attached and floating ribbons contained high levels of ^15^N. In fact, the majority of the ribbons showed a higher turnover than all other cellular components we analyzed (e.g., cytosol, nuclei or mitochondria; Fig. 7C, D). Several aged ribbons could be found exclusively in the membrane-attached population when investigating animals fed with the isotopic diet for 14 days; however, such ribbons were no longer identifiable after 21 days of feeding. Thus, our data suggest that: (i) VHC ribbons exhibit considerable turnover dynamics exceeding the ones observed for other organelles, (ii) different ribbons show different levels of turnover, even in the same cell – with a full exchange taking place every 1-3 weeks and importantly (iii) that floating ribbon clusters might be exclusively composed of newly formed ribbons, therefore pointing to normal formation and transport, but a faulty AZ membrane-attachment mechanism with advancing age.

## Discussion

The present study describes several morphological alterations in aging utricular VHCs (Fig. 8). Specifically, we found that: (i) both, VHC ribbons and mitochondria (mainly in type I VHCs), undergo drastic changes in shape and increase in size/volume during the aging process. (ii) Ribbons, successively, are found detached from the AZs and these floating ribbons start to accumulate, particularly in type I VHCs, in an age-dependent manner, thereby causing a remarkable increase in the total amount of ribbon material per cell. (iii) These floating ribbons are decorated with full sets of seemingly mature SVs and form clusters within the cytoplasm, often in proximity to the AZ. (iv) Ribbons within these clusters are mainly composed of newly-synthesized proteins. Finally, (v) our data suggests that these ‘new’ ribbons might collectively fail at the final membrane-anchoring step, potentially due to an age-dependent AZ depletion of the ribbon-anchoring protein bassoon.

**Fig. 8:**
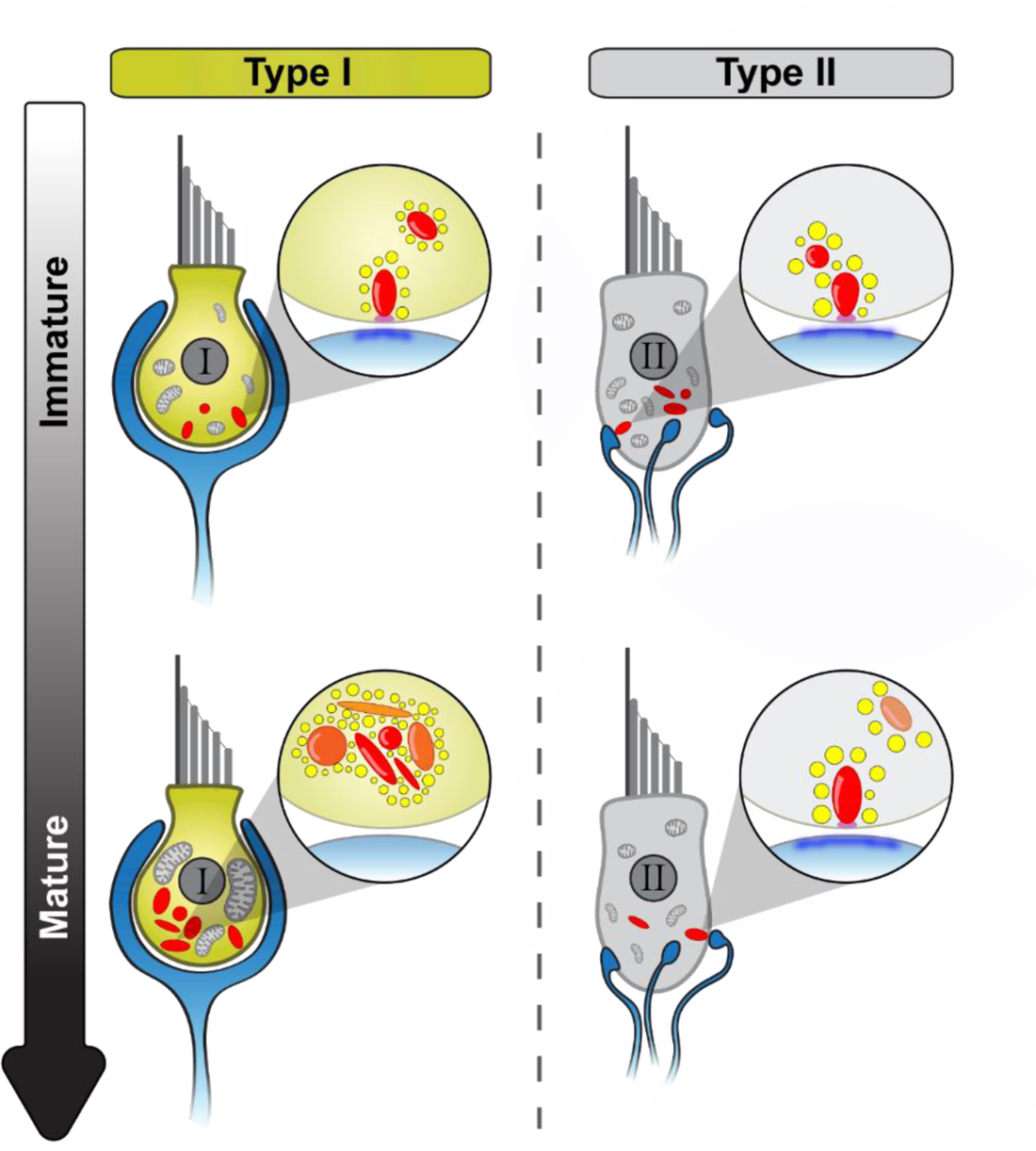
Schematic overview of utricular VHC ribbon synapse maturation and with advancing age. Summary of the main morphological observations during utricle development. While type II VHCs show very similar developmental events as detected in cochlear IHCs, type I VHCs undergo a distinct maturation process. Light blue: afferent contacts, red: synaptic ribbons (young/newly formed ribbons: transparent, old ribbons: opaque), yellow: SVs, magenta: presynaptic density, dark blue: PSD, light gray: mitochondria.

### VHC ribbon synapses undergo structural alterations in an age- and cell-type-dependent manner

First and foremost, this study reveals striking differences in the ribbons decorating type I and type II VHCs, and even more so when compared to the ribbons in closely-related cochlear IHCs. For example, our most prominent finding is a progressive accumulation of ‘floating’, i.e., not membrane-attached ribbons that, instead of being degraded, form large clusters in type I VHCs. While previous studies already reported on the occurrence of such cytosolic ribbon aggregates – for example, in VHCs of the murine *crista ampullaris*, ribbon clusters could be observed as early as P2-4 (Nordemar, 1983). Our study provides the first detailed longitudinal analysis of this phenomenon ranging from immature to late adult states.

Our observations in the maturing and older VHCs, approximately one year of age, stand in stark contrast to the developmental maturation of auditory IHCs, in which floating ribbons seem to present a characteristic, but transient, feature of cellular immaturity that disappears prior to hearing onset (Michanski et al., 2019; Sobkowicz et al., 1986; Wong et al., 2014). Consistent with our findings, floating ribbons were previously observed solely in adult type I VHCs, here specifically in female C57BL/6N mice; however, overall still more single ribbons than ribbon clusters were described (Park et al., 1987; Woods and Park, 1987). We assume that the ribbon number per cell and the ribbon number per cluster might have been underestimated due to the limitation of 2D data sets. The majority of estimates in our study, are thus based on the 3D volume reconstructions, which provide more accurate numbers of single ribbons as well as ribbons within clusters. At the age of 8 months, we found an average of approximately 2.5 single ribbons and ∼11 ribbons within clusters in type I VHCs. In type II VHCs at the same age 15 ribbons were single and 12.5 organized in clusters. The latter were smaller, containing a maximum of 4 ribbons, whereas up to 14 ribbons were found in a single cluster in type I VHCs. Interestingly, ribbon accumulations and floating ribbons were also detected in type I VHCs from a human fetus (Favre et al., 1986; Sans and Scarfone, 1996), indicating a similar arrangement in humans.

### Morphological similarities and differences between type I and type II VHC ribbons

While ribbons in both VHC types are of various sizes and shapes, an age-dependent transformation towards elongated ribbons was observed in type I VHCs only. In contrast, ribbon shape distributions in type II VHCs remained much more uniform across ages. Both cell types mostly anchor one to two ribbons per AZ and in both, the number of membrane-attached ribbons per AZ peaks at P15, with a small proportion of AZs involving three or four membrane-attached ribbons, in type II and I VHCs, respectively. Our results in type II VHCs are in line with a previous study in the vestibular end organs of adult guinea pigs showing that the AZs of type II VHCs mainly harbor single synaptic ribbons, while ∼20% of afferent terminals are opposed by two or more ribbons (Sjöbäck and Gulley, 1979).

A morphological characteristic that seems to be conserved between distinct hair cell types is the presence of translucent core ribbons (Liberman, 1980; Michanski et al., 2019; Sobkowicz et al., 1982; Stamataki et al., 2006). While the previous studies suggested that these ‘fenestrated’ ribbons represent the final stages in the lifespan of a ribbon, we could not find any age-related differences in their occurrence. This is in line with the work of Park et al. who observed such ribbons in both VHC types in the saccule of young and old mice (Park et al., 1987; Woods and Park, 1987), however, without the knowledge of the actual “ribbon age”. Since translucent cores tend to occur in larger ribbons, we hypothesize that such morphological changes may appear due to RIBEYE packaging limitation. Future studies employing NanoSIMS analysis might clarify whether the age of ribbons is a defining factor in the occurrence of this phenomenon.

### Microfilaments may serve as a means of transport for VHC ribbons

Ribbons were shown to form in the cytoplasm via auto-aggregation of the main structural scaffold protein RIBEYE (Magupalli et al., 2008). These aggregates form SV-tethering ribbon precursors that are then transported to the AZ in a microtubule (MT) and kinesin-dependent process (Michanski et al., 2019). However, VHCs are dominated by the occurrence of large strands of dense microfilaments that align with the apico-basal axis of mature type I VHCs. These filaments were previously described in otocyst VHCs during early development and were interpreted to be involved in nucleus migration (Heywood et al., 1975). While the molecular nature of these microfilaments remains elusive, functionally they might be involved in transport of sub-cellular structures, as they occupy a large apico-basal volume in aged type I VHCs. Moreover, since microfilament bundles were often observed in close proximity to ribbon clusters, it is conceivable that they contribute to ribbon transport towards the AZs. Alternatively, they might solely provide structural stability to VHCs. Future studies will be required to establish the molecular identity and function of these microfilaments.

### Aging affects mitochondrial volumes of utricular type I VHCs

The production, transport, and maintenance of ribbons, together with continuous tonic synaptic transmission are energy-demanding processes. Age-dependent cellular alterations, which ultimately attenuate release efficiency, may lead to energetic imbalance at the synapse that needs to be compensated. Such a scenario may be supported by our findings of enlarged synaptic mitochondria observed in aging type I VHCs. Mitochondria power the cells by producing ATP (Harris et al., 2012), which is for example required to synthesize glutamate (Waagepetersen et al., 2003), fill SVs with neurotransmitter molecules, prime SVs for release at the AZ membrane (Forgac, 2007; Verhage and Sørensen, 2008; Yao and Bajjalieh, 2008) or fuel local translation (Rossoll and Bassell, 2019). Moreover, an increased production of mitochondrial ATP is required for exocytosis and the endocytosis-based recycling of SVs, which even causes depletion of ATP during heavy cycling (Rangaraju et al., 2014). Therefore, it is not surprising that mitochondria were described to play a key role in quantal synaptic transmission (Smith et al., 2016; Wong et al., 2019). In the present study, we found drastically enlarged mitochondria in older type I VHCs. A similar observation of an performance- and age-dependent mitochondrial volume increase was reported e.g., in the hippocampus (Cserép et al., 2018; Pathak et al., 2015; Smith et al., 2016) and calyx of Held (Thomas et al., 2019), where mitochondria were found to be larger after hearing onset. The size enlargement was interpreted to support higher firing rates in auditory information processing to meet the higher energy demands and, thus, was proposed to be functionally-coupled to the release mode (Thomas et al., 2019). On one hand, since fewer ribbon synapses are attached in older type I VHCs, these ribbons have to work harder to achieve the same task, which might require higher energy. On the other hand, the accumulation of relatively young synaptic ribbons, together with a large number of SVs, also indicates an increased local translation of ribbon material, another energy demanding process (Rossoll and Bassell, 2019). Finally, the mitochondria ultimately supply energy for maintaining ionic homeostasis during continuous depolarization and hyperpolarization of the hair cells, which may be essential for type I VHCs since the supporting cells have no (or limited) access to the synaptic cleft.

Notably, in several studies, both vestibular and auditory disorders were associated with mitochondrial dysfunction (Böttger and Schacht, 2013; Fischel-Ghodsian et al., 2004; Iwasaki et al., 2011; Kokotas et al., 2007; Requena et al., 2014), which might also cause enlarged mitochondria.

### The formation of ribbon clusters increases with age and is a striking hallmark of older type I VHCs

With increasing age, the proportion of membrane-attached ribbons steadily declines, however, the prominence of large ribbon clusters, particularly in type I VHCs, successively increases. Here, accumulations of more than 10 ribbons could be shown to incapacitate several hundreds of SVs within a single cluster. While clusters can also be found in type II VHCs, their size typically remains moderate, commonly not exceeding 2-4 ribbons.

In our study, both types of VHCs have floating ribbons throughout adulthood, but with different abundance. The proportion of floating ribbons in type II VHCs remains mostly constant throughout the investigated ages (starting from early second postnatal week) and decreases modestly by ∼1 year of age, but detached ribbons never fully disappear. It is important to note that in rodents new type II VHCs can be formed throughout adulthood, although the regenerating potential is moderate (Burns and Stone, 2017; Forge et al., 1993; Forge et al., 1998; Golub et al., 2012; Hicks et al., 2020; Kawamoto et al., 2009; Lin et al., 2011; Slowik and Bermingham-McDonogh, 2013). Therefore, we cannot completely rule out the possibility that type II VHCs containing floating ribbons in old animals may represent (at least in part) freshly-regenerated and hence immature hair cells. Future experiments will be required to clarify this issue. Such a scenario is unlikely to take place in type I VHCs. In these cells, floating ribbons represent the vast majority of the total ribbon population in older mice, which additionally argues against these ribbons being a signature of newly regenerated cells. A hallmark of type I VHCs is their use of two modes of synaptic transmission, a classical quantal and a nonconventional non-quantal mode of synaptic transmission (reviewed in (Eatock, 2018; Mukhopadhyay and Pangrsic, 2022)). To what degree one or the other mode of transmission may operate in individual cells, and possibly influence and support each other to enable transmission of vestibular information, is under investigation (Contini et al., 2020; Highstein et al., 2015; Songer and Eatock, 2013; Spaiardi et al., 2022). One might like to speculate that floating ribbons may be a signature of “low” demand for quantal synaptic transmission. If so, the questions to address in the future would be, whether their abundance in cells changes dynamically upon demand or varies in distinct cell subpopulations? While, due to different technical limitations, we could not assess the absolute numbers of floating and attached ribbons in a large number of individual VHCs, we so far found no indication of distinct subpopulations of type I VHCs with different abundance of floating vs. attached ribbons, which remains an interesting question to be revisited in the future.

### An age-dependent deficit in ribbon membrane-anchoring at the AZ gives rise to floating ribbon clusters

The high abundance of floating ribbons in adult VHCs is highly unusual for ribbon-type synapses; however, various scenarios can be envisioned that may explain this phenomenon: (i) in the process of synaptic degeneration, ribbons may detach from the AZ membrane prior to degradation and/or recycling, thus accumulating in the cytoplasm, (ii) the process of formation and attachment of ribbons may be activity-dependent and highly dynamic, with cytosolic clusters in AZ proximity serving as a ‘reserve pool’ of ribbons that may be recruited to, or retracted from, the plasma membrane upon a change in demand, thus rapidly adding or removing substantial amounts of release-ready SVs, or (iii) ‘new’ ribbons may be formed continuously, but with increasing age, lose the ability to attach adequately at the plasma membrane.

The first hypothesis may be supported by a recent study that reported a selective loss of synaptic ribbons in extrastriolar type I VHCs of old mice (Wan et al., 2019). This may possibly be associated with detachment and subsequent degradation of synaptic ribbons, but such a mechanism remains to be unambiguously demonstrated.

Regarding the second hypothesis, floating ribbons were found in mice that were brought to space (Ross, 1994; Ross, 2000), suggesting a loss of attached ribbons due to a reduction of synaptic activity in response to hypogravity. However, since floating ribbons as well as ribbon clusters in rodents and humans are also present under normal-gravitational conditions, this explanation appears rather unlikely. Moreover, a significant decline of evoked vestibular potentials could only be detected at an age of two years in mice (Wan et al., 2019). Alternatively, floating ribbons may act as a dynamic reserve pool to supplement existing (or replace end-of-lifetime) ribbons as well as SVs in case of changes in demand. In this context, our NanoSIMS data revealed that some ribbons within a given cluster were slightly older than others and ribbon were newly formed and possibly partially replaced within a time line of 1.3 weeks. However, ribbons were considerably younger compared to other subcellular structures such as mitochondria. The extent of structural plasticity of ribbons is only partly understood to date; yet, in other sensory systems, such as retinal photoreceptors or pinealocytes, ribbons were shown to adapt their shape in response to the cellular state of activity (Fuchs et al., 2013; Spiwoks-Becker et al., 2004; Vollrath and Seidel, 1989; Voorn and Vogl, 2020). Of particular interest here is the observation that photoreceptor ribbons apparently regulate their size by pinching off or fusing with SV-bearing precursor spheres upon changes in illumination (Spiwoks-Becker et al., 2004). As discussed above, the abundance of VHC ribbon synapses may change in hypogravity (Ross, 2000; Sultemeier et al., 2017). If floating ribbons in VHCs also act as a reserve pool for SVs, is questionable. Judging from the low amount of depolarization-evoked membrane capacitance change (reporting SV fusion with the plasma membrane) in young type I cells (Spaiardi et al., 2022; Vincent et al., 2014), it appears unlikely that hundreds of SVs were in place to support very high rates of exocytosis and replenishment.

Finally – and in our opinion most likely – the occurrence of ribbon clusters may be a result of an age-dependent degenerative processes that affect stable ribbon anchoring at the AZ membrane. Our NanoSIMS data revealed that floating ribbons in particular have a high protein turnover, thereby arguing in favor of the idea that these ribbons represent the accumulation of ‘new’ ribbons. In support of this hypothesis, our FIB-SEM data further indicate that throughout the VHC lifespan additional ribbon material is formed and the total amount steadily increases. Either, the protein synthesis of ribbon material is not controlled well enough anymore or, in combination with the apparent reduction of bassoon levels at the AZ, render a scenario most likely in which ribbon synthesis and transport remain intact, but anchoring at the release site fails. Since bassoon is a well-characterized scaffold essential for adequate ribbon attachment to the AZ membrane across different sensory systems (Dick et al., 2003; Khimich et al., 2005), its involvement in this process would be expected. While the exact molecular mechanisms remain to be determined, our work adds important novel insights into VHC ribbon synapse physiology and thus provides new directions for future studies

## Materials and Methods

### Animals

C57BL/6J, CBA/J and C57BL/6JRj (Janvier laboratory) wild-type mice of either sex between the postnatal day (P)9 and 11 months were deeply anesthetized with CO_2_ and sacrificed by decapitation or cervical dislocation for immediate dissection of the utricle. For a subset of experiments, a Cre-dependent tdTomato reporter strain (*Ai14*; #007908 Jackson Laboratory; (Madisen et al., 2010) was crossbred with a *Neurog1^CreERT2^* mouse line (#008529 Jackson Laboratory; (Koundakjian et al., 2007) to sparsely label random vestibular ganglion neurons. All experiments complied with national animal care guidelines and were approved by the University of Göttingen Board for Animal Welfare and the Animal Welfare Office of the State of Lower Saxony.

### Immunohistochemistry, confocal and super-resolution STED microscopy

The inner ears were extracted from C57BL/6J mice at the age of 2 and 32 weeks. Small aperture was created in the bone just above the utricle with forceps. The pigmented membrane was carefully cut open to expose the utricle underneath. The entire inner ear was then transferred to 4% formaldehyde in phosphate buffer saline (PBS: Sigma #P4417, Germany) on ice for about 45 min to allow fixation of the tissues. Next, the sample was washed in deionized water for up to 1 minute and then briefly (ca. 30 s) transferred to TBD-1 (Shandon TBD-1^TM^ Rapid Decalcifier, Ref# 6764001) for decalcification, which was followed by another brief wash in deionized water at room temperature. The fixed and decalcified utricle was then carefully extracted from the inner ear using fine forceps and transferred to PBS.

The utricles were washed 3 times in PBS for 10 minutes each, followed by blocking with Goat Serum Dilution Buffer (GSDB: 16% normal goat serum, 450 mM NaCl, 0.3% Triton X-100, and 20 mM phosphate buffer, at pH 7.4) for 1 hour at room temperature in a wet chamber. The organs were then incubated overnight in a primary antibody concoction prepared in GSDB at 4-8 °C in a wet chamber. The following day the organs were washed 3 times in wash buffer for 10 minutes each and then incubated in a secondary antibody dilution in GSDB for 1 hour in a light protected wet chamber at room temperature. Finally, they were washed again 3 times in wash buffer before being mounted on microscopy slides with mounting medium containing Mowiol 4–88 and DABCO (Carl Roth, Germany) or ProLong Glass antifade mounting medium (ThermoFisher, Germany). The following antibodies and dilutions were used: chicken-anti-calretinin (Synaptic Systems, Germany, dilution 1:200), rabbit-anti-piccolino (Synaptic Systems, Germany, dilution 1:200), rabbit-anti-RIBEYE (Synaptic Systems, Germany, dilution 1:200), mouse-anti-bassoon (Abcam, Germany, dilution 1:300), mouse-anti-CtBP2 (BD Biosciences, USA, dilution 1:200) and rabbit anti-dsRed (Clontech, dilution 1:200) as well as a directly Atto-565 conjugated anti-RFP nanobody (NanoTag, dilution 1:100). The secondary antibodies used were: Alexafluor488-labeled goat-anti-chicken (ThermoFisher Scientific, USA, dilution 1:200), Alexafluor594 labeled goat-anti-mouse (ThermoFisher Scientific, USA, dilution 1:200) or STAR635p-labeled cameloid anti-mouse IgG1 nanobodies (NanoTag, dilution 1:100), as well as STAR580-labeled cameloid anti-rabbit nanobodies (NanoTag, dilution 1:100) or STAR635p-labeled goat-anti-rabbit (Abberior, Germany, dilution 1:200). Confocal and high resolution 2D-STED images were acquired using an Abberior Instruments Expert Line STED microscope equipped with 488, 561 and 640 nm excitation lasers, STED laser at 775 nm (1.2 W), and a 100x oil immersion objective (1.4 NA, Olympus). Images were acquired using the same microscope settings, analysed and processed in Fiji/ImageJ software (Schindelin et al., 2012) or Bitplane Imaris 9.6.1 (Oxford Instruments), and assembled in Affinity Designer 1.10.5 for display.

### Conventional embedding and transmission electron microscopy

Utricle organs from mice ranging in age between P9 to 11 months were isolated in HEPES Hank’s solution, which contained (in mM): 5.36 KCl (746436, Sigma, Germany), 141.7 NaCl (746398, Sigma, Germany), 10 HEPES (H3375, 006K5424, Sigma, Germany), 0.5 MgSO_4_-7H_2_O (Sigma, Germany), 1 MgCl_2_ (M2670, Sigma, Germany), 2 mg/ml D-glucose (G8270-1KG, Sigma, Germany), and 0.5 mg/ml L-glutamine (G3126-100G, #SLBS8600, Sigma, Germany) and was adjusted to pH 7.2, ∼300 mmol/kg. First the inner ear was dissected and the vestibular bone tissue was carefully removed. The inner ear was afterwards placed into a petri dish filled with proteinase XXIV solution (P 8038-100MG, Th. Geyer, Germany; 50 mg/ml in HEPES-HANKS solution, w/v) for 5 min at room temperature to facilitate the removal of the otolithic membrane overlying hair bundles of the utricle. In fresh HEPES Hank’s solution, the otolithic membrane was then removed and the exposed utricle was excised.

Conventional embeddings of vestibular organs were performed according to our previous studies (Michanski et al., 2019; Strenzke et al., 2016; Wong et al., 2014). In brief, the organs were fixed immediately after dissection with 4% paraformaldehyde (0335.1, Carl Roth, Germany) and 0.5% glutaraldehyde (G7651, Sigma, Germany) in PBS (P4417, Sigma, Germany; pH 7.4) for 1 h on ice followed by a second fixation step overnight with 2% glutaraldehyde in 0.1 M sodium cacodylate buffer (v/v, pH 7.2) at 4°C. Next, specimens were washed in 0.1 M sodium cacodylate buffer and treated with 1% osmium tetroxide (75632.5ml, Sigma, Germany; v/v in 0.1 M sodium cacodylate buffer) for 1 h on ice followed by further sodium cacodylate buffer and distilled water washing steps. After the *en bloc* staining with 1% uranyl acetate (8473, Merck, Germany; v/v in distilled water) for 1 h on ice, samples were briefly washed in distilled water, dehydrated in an ascending concentration series of ethanol, infiltrated and embedded in epoxy resin (R1140, AGAR-100, Plano, Germany) to get finally polymerized for 48 h at 70°C. Subsequently, ultrathin sections (70-75 nm) from the cured resin blocks were cut with an Ultracut E microtome (Leica Microsystems, Germany) or an UC7 microtome (Leica Microsystems, Germany) equipped with a 35° diamond knife (Diatome AG, Biel, Switzerland) and mounted on 1% formvar-coated (w/v in water-free chloroform) copper slot grids (3.05 mm Ø, 1 mm x 2 mm; Plano, Germany). Before imaging these sections at 80 kV using a JEM1011 transmission electron microscope (JEOL, Japan), they were counterstained with uranyl acetate and Reynold’s lead citrate (Reynolds, 1963) or uranyl acetate replacement solution (EMS, Science Services, USA). Micrographs of both hair cell types from the striolar and extrastriolar region were acquired at 10,000-x magnification with a Gatan Orius 1200A camera (Gatan GmbH, using the Digital Micrograph software package, USA).

### Electron tomography

Electron tomography was utilized as described previously in Strenzke *et al*. (Strenzke et al., 2016). Semithin sections (250 nm) were cut from conventionally embedded vestibular organs on an Ultracut E or an UC7 ultramicrotome (Leica Microsystems, Germany) with a 35° diamond knife (Diatome AG, Biel, Switzerland). Sections were placed on 1% formvar-coated (w/v in water-free chloroform) copper 100 mesh grids (3.05 mm Ø; Plano, Germany) and counterstained as described above. 10 nm gold beads (British Bio Cell, UK) were applied to both sides of the grids functioning as fiducial markers. With the Serial-EM software (Mastronarde, 2005), tilt series image acquisition was conducted mainly from −60 to +60° with 1° increments at 10,000-x magnification and a pixel size of 1.43 nm using a JEM2100 (JEOL, Japan) transmission electron microscope at 200 kV. For final tomogram alignments, the IMOD software package etomo was used and tomographic reconstructions were generated using 3dmod (Kremer et al., 1996). Movies were generated with IMOD, Fiji (Schindelin et al., 2012) and Windows Movie Maker 2012.

### Focused ion beam (FIB)-scanning electron microscopy (SEM)

Enhanced *en bloc* staining for FIB-SEM samples was performed according to Deerinck *et al*. (Deerinck et al., 2018). After fixation (similar to the above-described conventional embedding protocol), utricular organs were treated with a 1.5% potassium ferrocyanide (EMS, USA) and 4% osmium tetroxide solution (v/v in 0.1 M sodium cacodylate buffer) for 1 h on ice. Next, specimens were briefly washed in distilled water and placed in a thiocarbohydrazide (w/v in distilled water) solution for 20 min followed by additional washing steps in distilled water. A second exposure to 2% osmium tetroxide (v/v in 0.1 M sodium cacodylate buffer) was conducted followed by brief washing steps in distilled water before the samples were placed in 2.5% uranyl acetate (v/v in distilled water) overnight at dark. Subsequently, samples were washed in distilled water and contrasted with Reynold’s lead citrate (Reynolds, 1963) for 30 min at 60°C to be finally washed once again in distilled water, dehydrated in increasing ethanol concentrations, infiltrated and embedded in Durcupan (25%, 50%, 75% Durcupan in acetone for 1 h each and 100% Durcupan overnight; 44610, Sigma, Germany) to get polymerized for 48 h at 60°C. After trimming the cured blocks with a 90° diamond trimming knife (Diatome AG, Biel, Switzerland), the blocks were attached to SEM stubs (Science Services GmbH, Pin 12.7 mm x 3.1 mm) with a silver filled epoxy (Epoxy Conductive Adhesive, EPO-TEK EE 129-4; EMS, USA) and polymerized at 60° overnight. Samples were coated with a 10 nm platinum layer using the sputter coating machine EMACE600 (Leica, Germany) at 30 mA current to be finally placed into the Crossbeam 540 focused ion beam scanning electron microscope (Carl Zeiss Microscopy GmbH, Germany) and positioned at an angle of 54°. A 400 nm platinum layer was deposited on top of the regions of interest and the Atlas 3D (Atlas 5.1, Fibics, Canada) software was used to collect the 3D data. Specimens were exposed to the ion beam driven with a 15 or 30 nA current while a 7 or 15 nA current was applied to polish the cross-section. Images from both hair cell types of the striolar and extrastriolar region were acquired at 1.5 kV using the ESB detector (450 or 1500 V ESB grid, pixel size x/y 3 or 5 nm) in a continuous mill and acquire mode using 700 pA or 1.5 nA for the milling aperture (z-step 5 nm). For subsequent post processing, data were aligned using the Plugin “Linear Stack Alignment with SIFT”, inverted and cropped in Fiji. Depending on the dataset properties, a smoothing function (3×3), local contrast enhancement using a CLAHE plugin in Fiji, and a binning by 2 in x/y was applied (Schindelin et al., 2012). Movies were generated with IMOD (Kremer et al., 1996), Fiji, Blender (www.blender.org) and Windows Movie Maker 2012.

### Nanoscale secondary ion mass spectrometry (NanoSIMS)

Mice were habituated to unlabeled ^14^N-SILAM diet for 1 week before labeling (Silantes, Germany; cat. Num. 231,004,650). 10 and 14 months old C57BL/6JRj mice were then labeled with the ^15^N-SILAM diet (Silantes, Germany; cat. Num. 231,304,650), for 14 or 21 days before the inner ear perfusion with a 4% paraformaldehyde (0335.1, Carl Roth, Germany) and 0.5% glutaraldehyde (G7651, Sigma, Germany) in PBS (P4417, Sigma, Germany; pH 7.4) fixative for 1 h on ice. Afterwards, inner ears were placed in a second fixative solution overnight at 4°C (2% glutaraldehyde in 0.1 M sodium cacodylate buffer (v/v, pH 7.2)). On the second day fixed utricular organs were excised and the conventional embedding protocol was performed as described earlier in detail. The epoxy resin (R1140, AGAR-100, Plano, Germany) embedded samples were, similar to other conventional embedded specimens, further processed for transmission electron microscopy. 110 nm sections were cut using a 35° diamond knife (Diatome AG, Biel, Switzerland) with an UC7 ultramicrotome (Leica Microsystems, Germany) and placed on 200-mesh copper finder grids (01910-F, Ted Pella). Regions of interest (ROIs) were selected with a JEM1011 (JEOL, Japan) at 80 kV using different magnifications (x80, x150x, x800, x2,500, x6,000, x8,000). Subsequently, NanoSIMS imaging was performed. The areas of interest, previously imaged by TEM, were imaged with a NanoSIMS 50 L (Cameca, Gennevilliers Cedex, France) using an 8 kV Cs^+133^ primary ion source. To reach the steady-state of the secondary ion yield, prior to each measurement, an area of 100×100 µm was implanted applying a current of 50 pA for ∼3 min (primary aperture D1:1). Subsequently, a primary ion current of ∼0.25 pA (primary aperture D1:5) was applied to obtain images of 256 x 256 pixels from areas of 10 x 10 or 5 x 5 µm, resulting in a pixel size of 39.06 or 19.53 nm respectively. The dwell time was 5 ms per pixel and three consecutive layers were accumulated for each final image. The detectors were set to collect the following ions: ^12^C^14^N^−^, ^12^C^15^N^−^, ^31^P^-^ and ^32^S^-^, and the mass resolving power of the instrument was adjusted to ensure the discrimination between isobaric interferences such as ^12^C^15^N^−^ from ^13^C^14^N^−^, or ^12^C^14^N^−^ from ^12^C_2_ H_2_ by using an entrance slit of 20×140µm and an aperture slit of 200 x 200 µm. The images were then exported and processed using WinImage (Cameca, Gennevilliers Cedex, France). The following masses were collected for each run: ^12^C^14^N (referred to as ^14^N in this report), ^12^C^15^N (referred to as ^15^N in this report), and ^31^P. ^31^P peak was used to mark the location of cellular structures. Each image shown in this manuscript is the result of a summation of all three image layers taken during analysis.

### Data analysis and statistics

<u>Quantitative analysis of electron microscopic random 2D sections</u> was performed with ImageJ/Fiji (Schindelin et al., 2012) as follows:

In order to quantify the occurrence of floating ribbons, we defined “floating ribbons” by a lack of any physical contact to the presynaptic density, while an “attached ribbon” exhibited a clear membrane anchorage via a presynaptic density. For the ribbon size of attached ribbon synapses, the height and width were measured taking the longest axis of the ribbon excluding the presynaptic density as well as manual tracing of the synaptic ribbon was performed in order to determine the ribbon area. In contrast, due to the lack of an attachment of floating ribbons to the membrane, no precise ribbon orientation could be identified and hence the size of floating ribbons could only be analyzed by measuring the ribbon area.

For SVs of attached and floating ribbons, first the total amount of vesicles ≤ 80 nm from the ribbon surface were quantified in number and size. The latter analysis included the horizontal and vertical axis measurements, which were averaged to calculate the mean SV diameter. Next, the SV density was calculated by dividing the total number of SVs by the ribbon surface area.

Exclusively for attached ribbon synapses, two distinct morphological vesicle pools were additionally quantified in terms of numbers (as previously characterized in (Strenzke et al., 2016)): (i) membrane-proximal synaptic vesicles (MP-SVs, ≤ 25 nm distance between SV membrane and AZ membrane and ≤ 80 nm from the presynaptic density); and (ii) ribbon-associated synaptic vesicles (RA-SVs, first layer of vesicles around the ribbon with a maximum distance of 80 nm from the ribbon surface to the vesicle membrane and not falling into the MP-SV pool).

#### Quantitative analysis of FIB-SEM data

FIB-SEM datasets were segmented semi-automatically using the 3dmod package from the IMOD software (Kremer et al., 1996) and applying the imodinfo function, information about the ribbon and mitochondria volume was given. Distance measurements were performed with the measurement drawing tool along the x, y and z-axis.

#### Quantitative analysis of NanoSIMS data

NanoSIMS images were analyzed using a custom-written Matlab macro (the Mathworks Inc., Natick, MA, USA). NanoSIMS images were overlaid with the EM images, using Adobe Photoshop CS6 (Adobe Systems Incorporated). ROIs were selected manually on the EM images, and the ^15^N and ^14^N counts were measured for all ROIs in the respective NanoSIMS images. The results were saved as text files and were then combined to provide the statistics shown in the NanoSIMS figure.

All data are mainly presented as boxplots with individual data points overlaid and as bar graphs highlighting the mean or percentages of the data, which were analyzed using Excel, Igor Pro 6 (Wavemetrics Inc., USA), R and Java, if not otherwise indicated. Normality was assessed with the Jarque-Bera test and equality of variances in normally distributed data was assessed with the F-test. In order to compare two samples, the two-tailed unpaired Student’s t-test, or, when data were not normally distributed and/or variance was unequal between samples, the unpaired two-tailed Mann-Whitney-Wilcoxon test was used. One-way ANOVA test (followed by Fischer post-hoc tests (NanoSIMS data)) followed by Tukey’s test was used to calculate statistical significance in multiple comparisons for normally distributed data or in case of non-normally distributed data the Kruskal-Wallis (KW) test followed by non-parametric Employing the analysis tool based on Java Statistical Classes library (JSC) (Bertie, 2002) applied in our previous study (Jean et al., 2018), we performed the KW test for the identification of significant differences between SV diameters. Non-significant differences between samples are indicated as *n.s*., significant differences are indicated as *\*P* < 0.05, *\*\*P* < 0.01, *\*\*\*P* < 0.001, *\*\*\*\*P* < 0.0001. Affinity Designer 1.10.5 and Adobe Illustrator 2023 (Adobe Inc., USA) were utilized for Figure assembling and display.

## Acknowledgements

We thank S. Gerke, C. Senger-Freitag, A.J. Goldak, S. Langer, I. Preuss, C. Fischer, A. Zeise and C. Förster for expert technical assistance, O. L. Diaz for IT support. This work was funded by German Research Foundation grants: Collaborative Research Center 889 [Projects A07 (C.W.), B08 to C.V., and B09 to T.P.] and T.P., S.J. and C.W. were further supported by the Deutsche Forschungsgemeinschaft under Germany’s Excellence Strategy - EXC 2067/1-390729940. S.O.R., S.J. and C.W. were further funded by German Research Foundation grants: Collaborative Research Center 1286 [Projects A04 to C.W., A05 to S.J., and A03 to S.O.R.]. Part of this work (A.M.S.) was funded by the Cluster of Excellence and DFG Research Center Nanoscale Microscopy and Molecular Physiology of the Brain [Research field A1 (W.M.)] and by the Deutsche Forschungsgemeinschaft (DFG) (FOR2848, MO 1082/1-2, project 08 to W.M.). C.V. was an Otto Creutzfeldt-Fellow of the Elisabeth and Helmut Uhl Foundation. M.M. and T.P. are funded via the BMBF and MWK (Bundesministerium für Bildung und Forschung and Niedersächsisches Ministerium für Wissenschaft und Kultur; Professorinnenprogramm III, 22-38 285/1-3-12-1 and 22 - 76251-99-17/19 to T.P.). E.F.F. was supported by a Schram Stiftung (T0287/35359/2020) and a DFG grant (FO 1342/1-3). S.O.R., K.G. and P.G.A. were supported by a DFG grant (RI 1967/10-1, NeuroNex).

## Author Contributions

S.M. and C.W. designed the study. S.M. performed electron microscopic work (conventional embeddings, enhanced en bloc stainings, TEM of random sections, electron tomography, FIB-SEM). T.H. contributed to 2D data analysis and acquisition. T.P., M.M. and C.V. performed immunohistochemistry as well as confocal microscopy and STED. S.M., M.G., T.H., M.M. and C.V. analyzed data. A.M.S. performed and W.M. supervised FIB-SEM. M.G. contributed to statistical analysis. P.I. made the Blender movies. S.J. and P.I. contributed to mitochondria data interpretation. E.F.F. prepared mice for the NanoSIMS experiments. S.M. and E.F.F. prepared tissue samples for NanoSIMS. C.W. performed TEM image acquisitions for NanoSIMS. P.A.G. and K.G. performed NanoSIMS and S.O.R. and P.A.G. analyzed NanoSIMS data. S.M. and C.W. prepared the manuscript with contributions from all other authors.

---

**Fig. S1:**
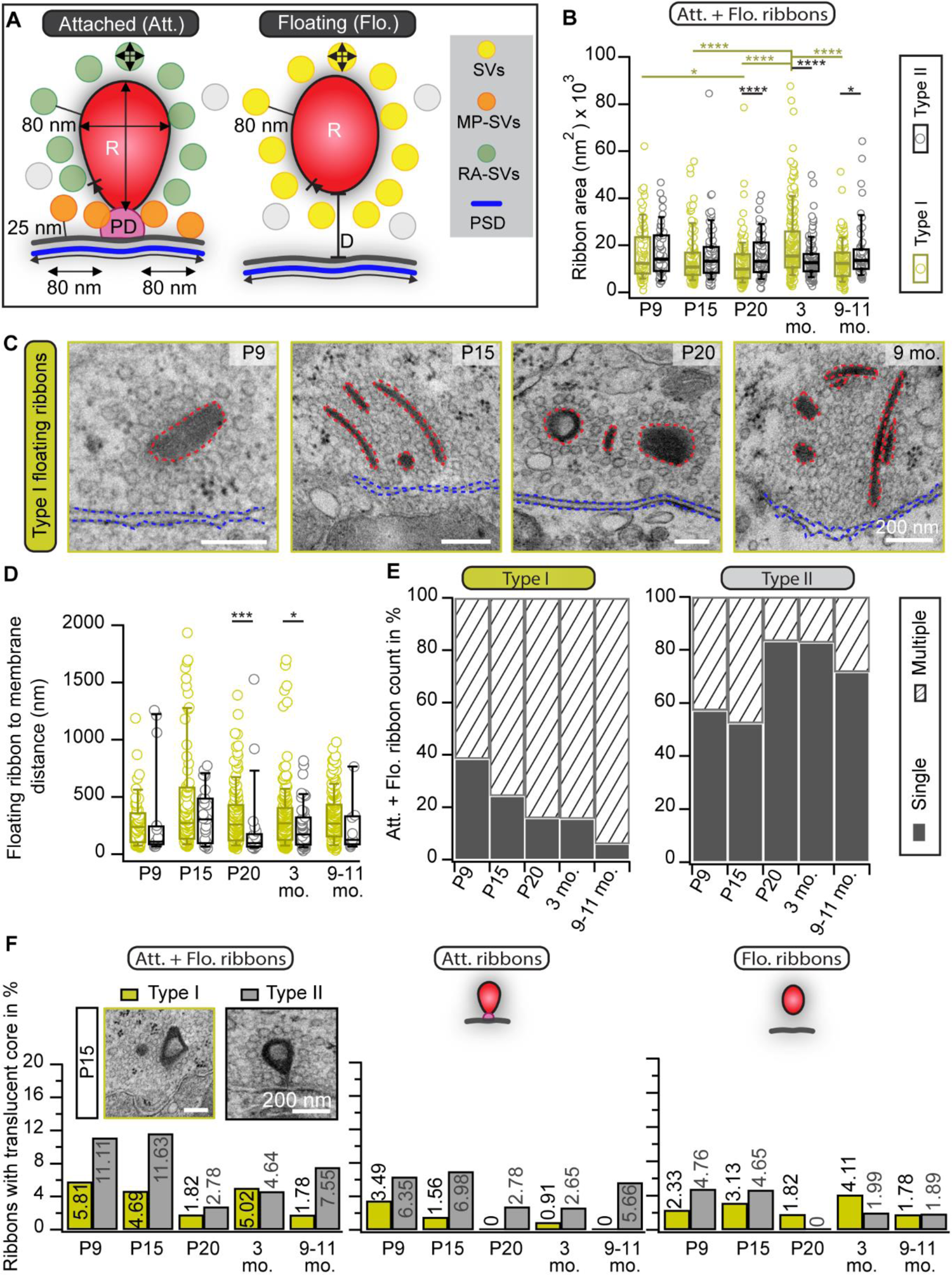
With advancing age the number of multiple ribbons rises in type I VHCs but reduces in type II VHCs. (A) Schematic representation of the applied quantification criteria. (B) Box plot of ribbon area measurements from both attached (Att.) as well as floating (Flo.) ribbons, respectively for all age groups. (C) Representative TEM images displaying more examples of floating ribbons and ribbon clusters in type I VHCs. (D) Box plot with individual data points from the distance measurements of floating ribbons to the cell membrane. (E) The combination of ribbon counts from the two categories, attached and floating ribbons, shows a remarkable opposing frequency gradient of single and multiple ribbons between the VHC types. (F) Percentages of type I and II VHC ribbon synapses that exhibit a translucent core.

**Fig. S2:**
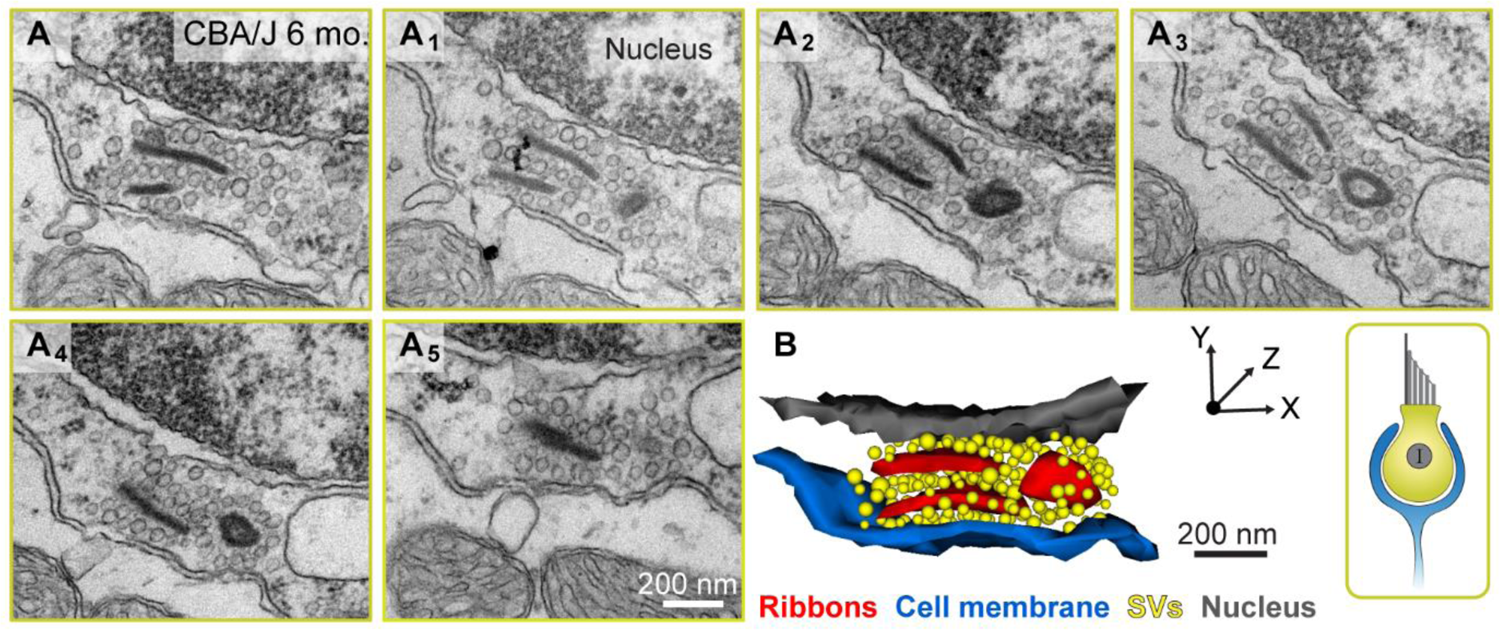
Floating ribbon clusters are also present in adult CBA/J mice. (A-A5) Consecutive ultrathin sections displaying a ribbon cluster of three adjacent floating ribbons and the corresponding 3D model (B).

**Fig. S3:**
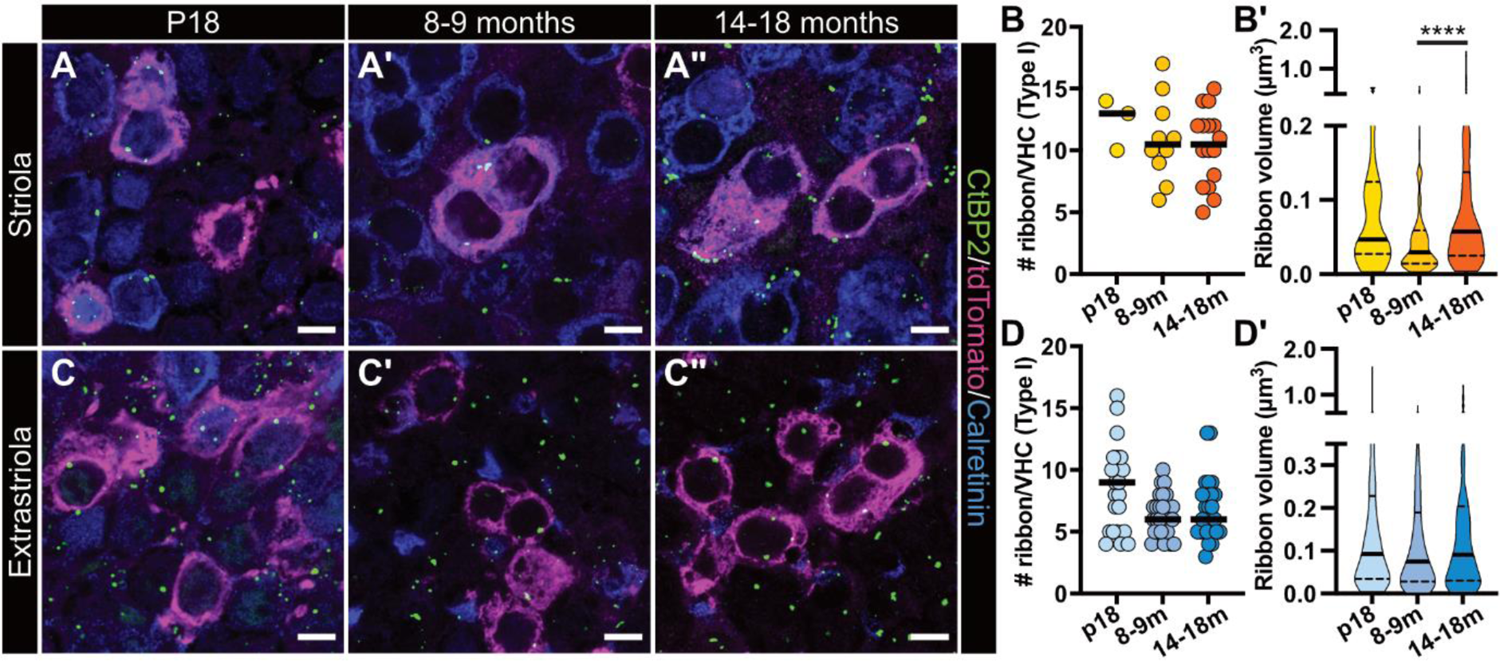
Larger synaptic ribbon spots in old type I VHCs in the striola. Representative confocal maximum projections of (A-A”) striolar and (C-C”) extrastriolar VHCs of a *Ai14-Neurog1-creER^T2^* knock-in mouse – that sparsely expresses tdTomato in a random subset of vestibular ganglion neurons – stained for the ribbon marker CtBP2 (green), tdTomato (magenta) and the Ca^2+^-buffer calretinin (blue). (B and D) Plotted are individual cells as well as (B’ and D’) individual ribbon volumes per age group and utricular location. In the striolar region, only VHCs with calretinin-positive calyces were considered for analysis, whereas in the extrastriolar region only VHCs with calretinin-negative calyces were analyzed. Ribbon counts in (B and D) consist of data from 3 striolar and 23 extrastriolar type I VHCs from a single P18 animal, while the 8-9 months group consists of 10 striolar and 28 extrastriolar type I VHCs from N=3 animals and the 1.5y group of 16 striolar and 29 extrastriolar type I VHCs from N=3 individuals. Ribbon volumes in (B’ and D’) are displayed with violin plots, where the thick line indicates the median and the thinner dashed lines the inner and outer quartiles. Data were derived from a total of 37, 108 and 174 striolar as well as 196, 180 and 194 ribbons of the respective age groups. *\*\*\*\*P*<0.0001 (KW test).

**Fig. S4:**
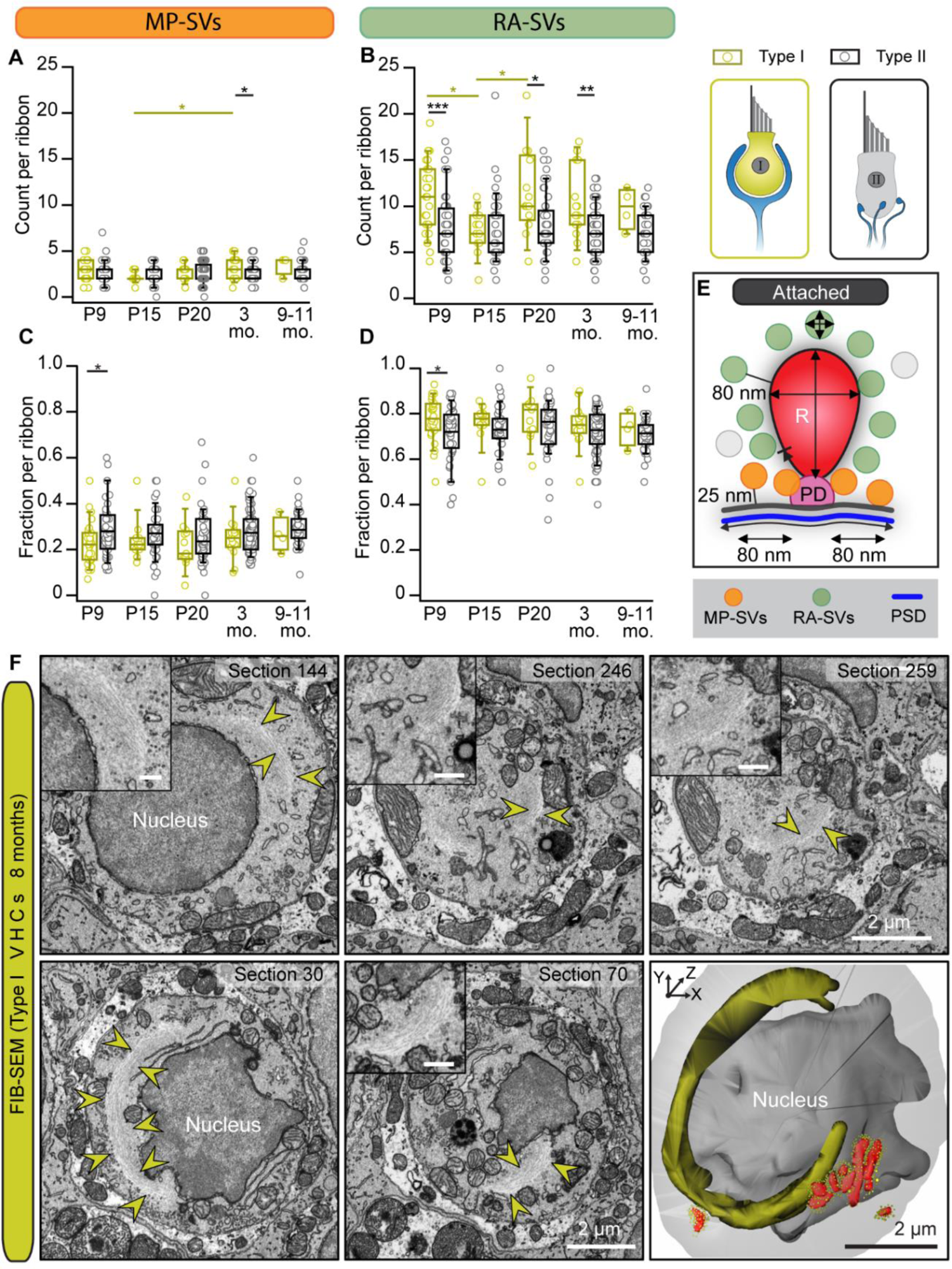
Extended strands of microfilaments close to ribbon clusters in mature type I VHCs. (A-D) SV count data and their fraction per ribbon for the two different SV pools of attached ribbons. (E) Schemata of the applied quantification criteria for MP-vs. RA-SV pool. (F) Cytoskeletal strands distributions were traced in 3D and are exemplarily highlighted (yellow arrowheads) in individual sections from two different FIB-SEM stacks. In the lower panel, the microfilaments (darker yellow in the 3D model) end close to a ribbon cluster (red objects in the 3D model) at the basal part of the type I VHC. For all insets the scale bars are 500 nm.

**Fig. S5:**
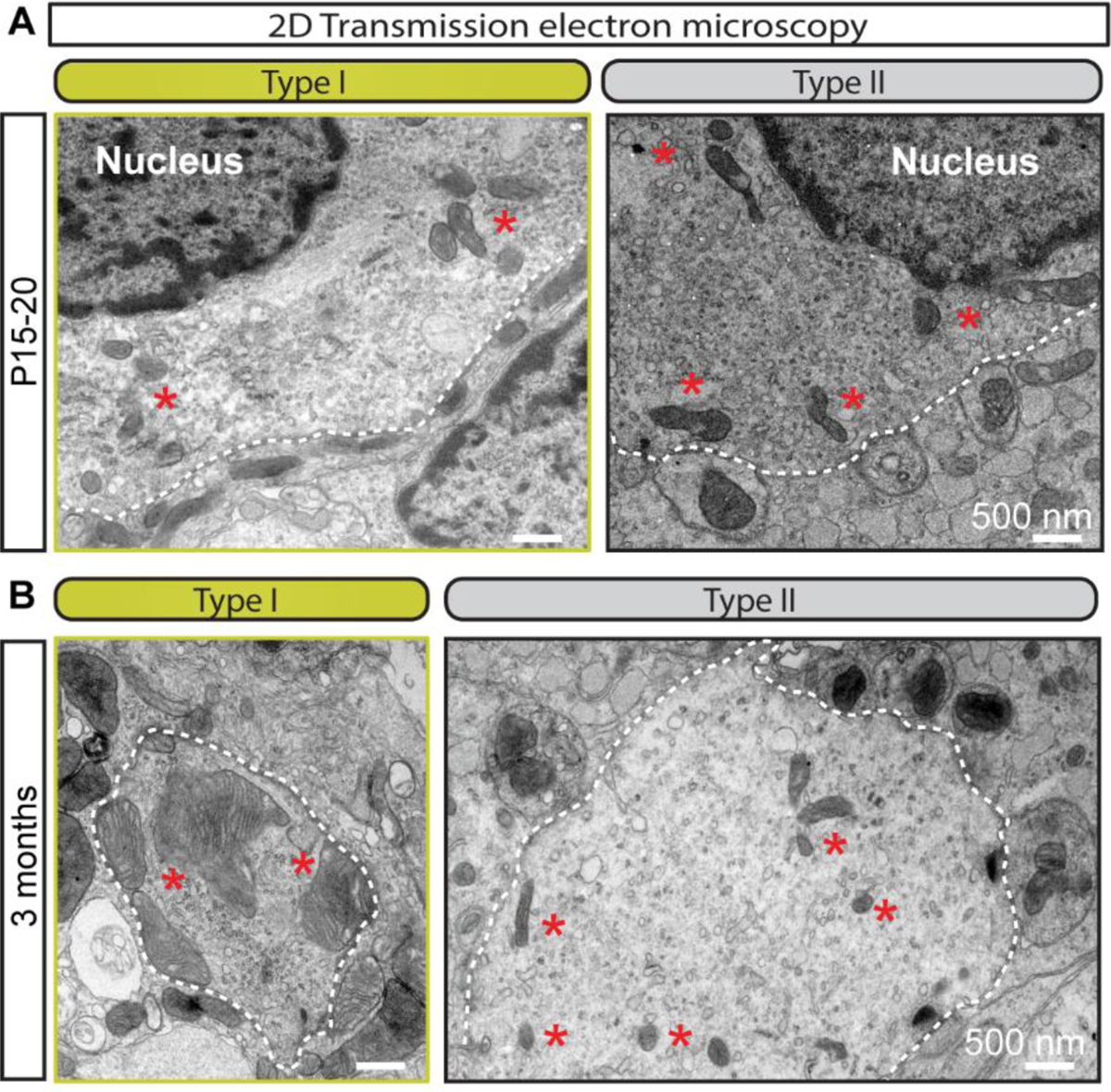
Mitochondrial size increases in type I VHCs with advancing age as visualized with 2D electron microscopy. Representative electron micrographs highlighting different sized mitochondria (red asterisks) from young ages (A) compared to a mature age (B).

**Table S1:**
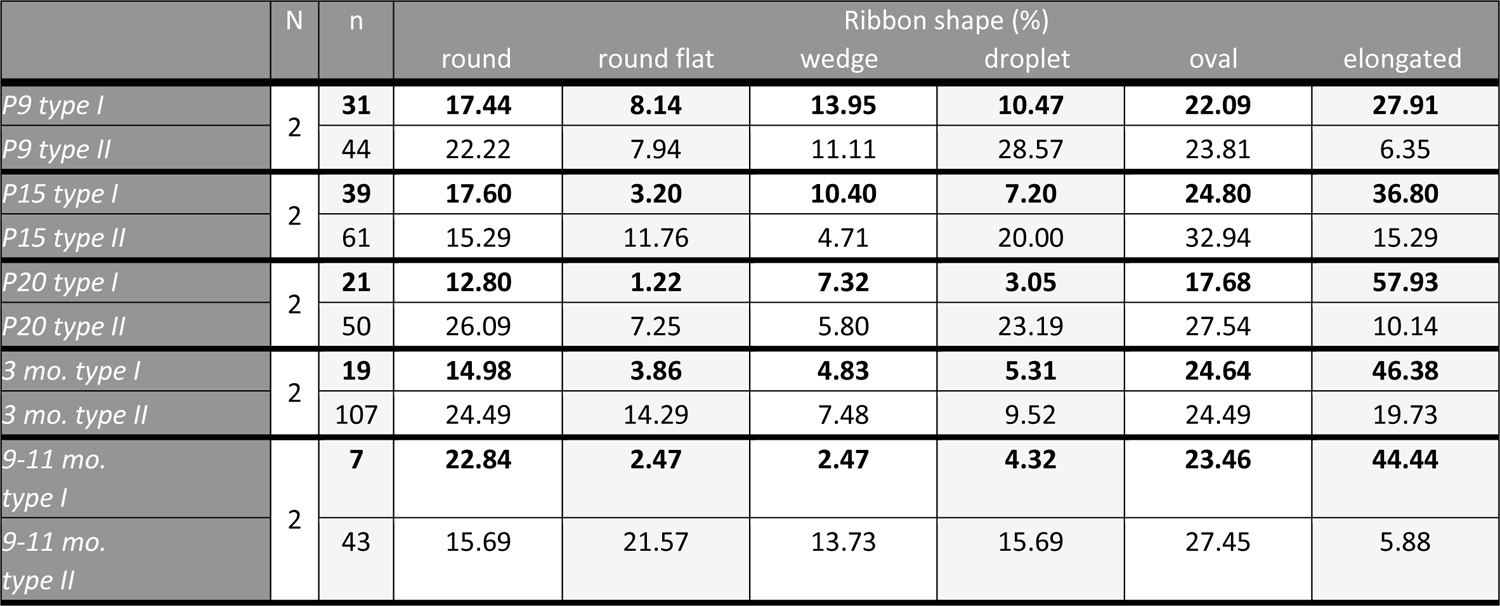
Evaluation of ribbon shapes observed by random section TEM. Percentages of ribbon shapes at type I and type II VHCs. N = animal count, n = ribbon count (Supplementary data for Fig. 1).

**Table S2:** Ribbon size measurement data from TEM. Attached (Att.) and floating (Flo.) ribbon size values from type I and type II VHCs (Supplementary data for Fig. 1 and Fig. S1). Data are presented as median ± SEM. *P*-values are calculated by the two-tailed unpaired Student’s t-test for normally distributed data with equal variances or the unpaired two-tailed Mann-Whitney-Wilcoxon test was applied for not normally distributed data and/or when variances were unequal between two samples. For multiple comparisons, the one-way ANOVA with post-hoc Tukey’s test was used for normally distributed data or in case of non-normally distributed data the KW test with multiple comparison correction was utilized. Significant results are highlighted with *\*P* < 0.05, *\*\*P* < 0.01, *\*\*\*P* < 0.001, *\*\*\*\*P* < 0.0001. N = animal count, n = ribbon count.

|  | N | n | Ribbon area (nm<sup>2</sup>) x 10<sup>3</sup><br>(Att. + Flo.) | Ribbon area (nm<sup>2</sup>) x 10<sup>3</sup><br>(Att.) | Ribbon area (nm<sup>2</sup>) x 10<sup>3</sup><br>(Flo.) | Ribbon height (nm)<br>(Att.) | Ribbon width (nm)<br>(Att.) |
| --- | --- | --- | --- | --- | --- | --- | --- |
| P9 type I | 2 | 31 | 12.33 ± 1.27 | 18.70 ± 2.55 | 10.58 ± 1.17 | 194.75 ± 11.33 | 125.12 ± 13.81 |
| P9 type II |  | 44 | 14.10 ± 1.31 | 13.86 ± 1.56 | 18.21 ± 2.43 | 157.81 ± 8.50 | 104.48 ± 7.03 |
| P15 type I | 2 | 39 | 10.76 ± 0.89 | 13.27 ± 1.32 | 10.7 ± 1.14 | 179.15 ± 10.54 | 93.21 ± 6.94 |
| P15 type II |  | 61 | 13.21 ± 1.25 | 11.51 ± 1.09 | 15.94 ± 3.34 | 145.99 ± 10.90 | 95.09 ± 5.40 |
| P20 type I | 2 | 21 | 9.89 ± 0.70 | 15.33 ± 3.40 | 9.50 ± 0.60 | 241.14 ± 22.56 | 67.82 ± 23.01 |
| P20 type II |  | 50 | 13.16 ± 1.05 | 13.32 ± 1.23 | 12.13 ± 2.04 | 162.29 ± 6.44 | 116.22 ± 7.22 |
| 3 mo. type I | 2 | 19 | 15.45 ± 0.97 | 16.58 ± 5.87 | 15.19 ± 0.82 | 181.16 ± 18.03 | 134.33 ± 16.86 |
| 3 mo. type II |  | 107 | 12.62 ± 0.61 | 12.49 ± 0.68 | 13.06 ± 1.32 | 149.02 ± 3.62 | 103.48 ± 4.13 |
| 9-11 mo. type I | 2 | 7 | 12.33 ± 0.60 | 17.57 ± 3.45 | 12.30 ± 0.60 | 161.70 ± 25.53 | 97.24 ± 31.00 |
| 9-11 mo. type II |  | 43 | 13.60 ± 1.67 | 13.63 ± 1.84 | 12.00 ± 4.81 | 148.48 ± 8.90 | 118.58 ± 7.79 |
| P-values |  |  | **\*\*\*\* 3 mo. type I vs. P15 type I**<br>**P20 type I**<br>**9-11 months type I**<br>3 mo. type II<br>**** P20 type I vs. type II<br>*P = 0.045<br>9-11 months type I vs. type II<br>*P = 0.0133 P9 type I vs. P20 type I | *P = 0.035<br>P9 type I vs. type II<br>***P = 0.0008<br>3 mo. type I vs. type II | **P = 0.009<br>P15 type I vs. type II<br>*P = 0.011<br>3 mo. type I vs. type II<br>**** 3 mo. type I vs. P15 type I<br>P20 type I<br>9-11 months type I<br>**P = 0.0033<br>3 mo. type I vs. P9 type I | **** P20 type I vs. type II<br>**P = 0.010<br>3 mo. type I vs. type II<br>*P = 0.027<br>P15 type I vs. type II<br>*P = 0.0425<br>P20 type I vs. P15 type I | *P = 0.018<br>P20 type I vs. type II<br>*P = 0.0279<br>P9 type I vs. P20 type I |

**Table S3:** TEM analysis for ribbon counts. Percentages of ribbon numbers at type I and type II VHCs. N = animal count, n = ribbon count (Supplementary data for Fig. 2 and Fig. S1).

|  | N | n | Ribbon count (%) |  | Ribbon count in (%) |  | Ribbon count per cluster (%) |  |  |  |  |  |  |  |
| --- | --- | --- | --- | --- | --- | --- | --- | --- | --- | --- | --- | --- | --- | --- |
|  |  |  | Attached | Floating | Single | Multiple | 2 | 3 | 4 | 5 | 6 | 7 | 8 | 9 |
| *P9 type I* | 2 | **86** | **36.05** | **63.95** | **38.37** | **61.63** | **40.00** | **33.33** | **13.33** | **6.67** | - | **6.67** | - | - |
| *P9 type II* |  | 63 | 69.84 | 30.16 | 57.14 | 42.86 | 80.00 | 20.00 | - | - | - | - | - | - |
| *P15 type I* | 2 | **128** | **30.47** | **69.53** | **24.22** | **75.78** | **40.00** | **16.00** | **20.00** | **20.00** | - | - | **4.00** | - |
| *P15 type II* |  | 86 | 70.93 | 29.07 | 52.33 | 47.67 | 44.44 | 33.33 | 22.22 | - | - | - | - | - |
| *P20 type I* | 2 | **165** | **12.73** | **87.27** | **15.76** | **84.24** | **34.15** | **29.27** | **21.95** | **4.88** | **7.32** | **2.44** | - | - |
| *P20 type II* |  | 72 | 69.44 | 30.56 | 83.33 | 16.67 | 100.00 | - | - | - | - | - | - | - |
| *3 mo. type I* | 2 | **219** | **8.68** | **91.32** | **15.53** | **84.47** | **33.93** | **30.36** | **23.21** | **1.79** | **8.93** | - | - | **1.79** |
| *3 mo. type II* |  | 151 | 70.86 | 29.14 | 82.78 | 17.22 | 77.78 | 22.22 | - | - | - | - | - | - |
| *9-11 mo. type I* | 2 | **169** | **4.14** | **95.86** | **5.92** | **94.08** | **16.22** | **21.62** | **24.32** | **10.81** | **16.22** | **8.11** | **2.70** | - |
| *9-11 mo. type II* |  | 53 | 81.13 | 18.87 | 71.70 | 28.30 | 50.00 | 50.00 | - | - | - | - | - | - |

**Table S4:** FIB-SEM data on ribbon count quantifications. Percentages of ribbon numbers at type I and type II VHCs. N = animal count, n = ribbon count (Supplementary data for Fig. 4).

|  | N | n | Ribbon count (%) |  | Ribbon count (%) |  | Ribbon count per synapse (%) |  |  |  |  |  |  |  |  |
| --- | --- | --- | --- | --- | --- | --- | --- | --- | --- | --- | --- | --- | --- | --- | --- |
|  |  |  | attached | floating | single | multiple | 1 | 2 | 3 | 4 | 5 | 8 | 9 | 12 | 14 |
| *P15 type I* | 2 | **37** | **51.35** | **48.65** | **29.73** | **70.27** | **64.71** | **17.65** | **5.88** | - | **5.88** | - | - | **5.88** | - |
| *P15 type II* | 1 | 26 | 96.15 | 3.85 | 15.38 | 84.62 | 45.45 | 27.27 | 9.09 | 9.09 | - | 9.09 | - | - | - |
| *8 mo. type I* | 2 | **53** | **13.21** | **86.79** | **18.87** | **81.13** | **62.50** | **6.25** | - | - | **12.50** | **6.25** | **6.25** | - | **6.25** |
| *8 mo. type II* |  | 55 | 94.55 | 5.45 | 54.55 | 45.45 | 78.05 | 12.20 | 7.32 | 2.44 | - | - | - | - | - |

**Table S5:**
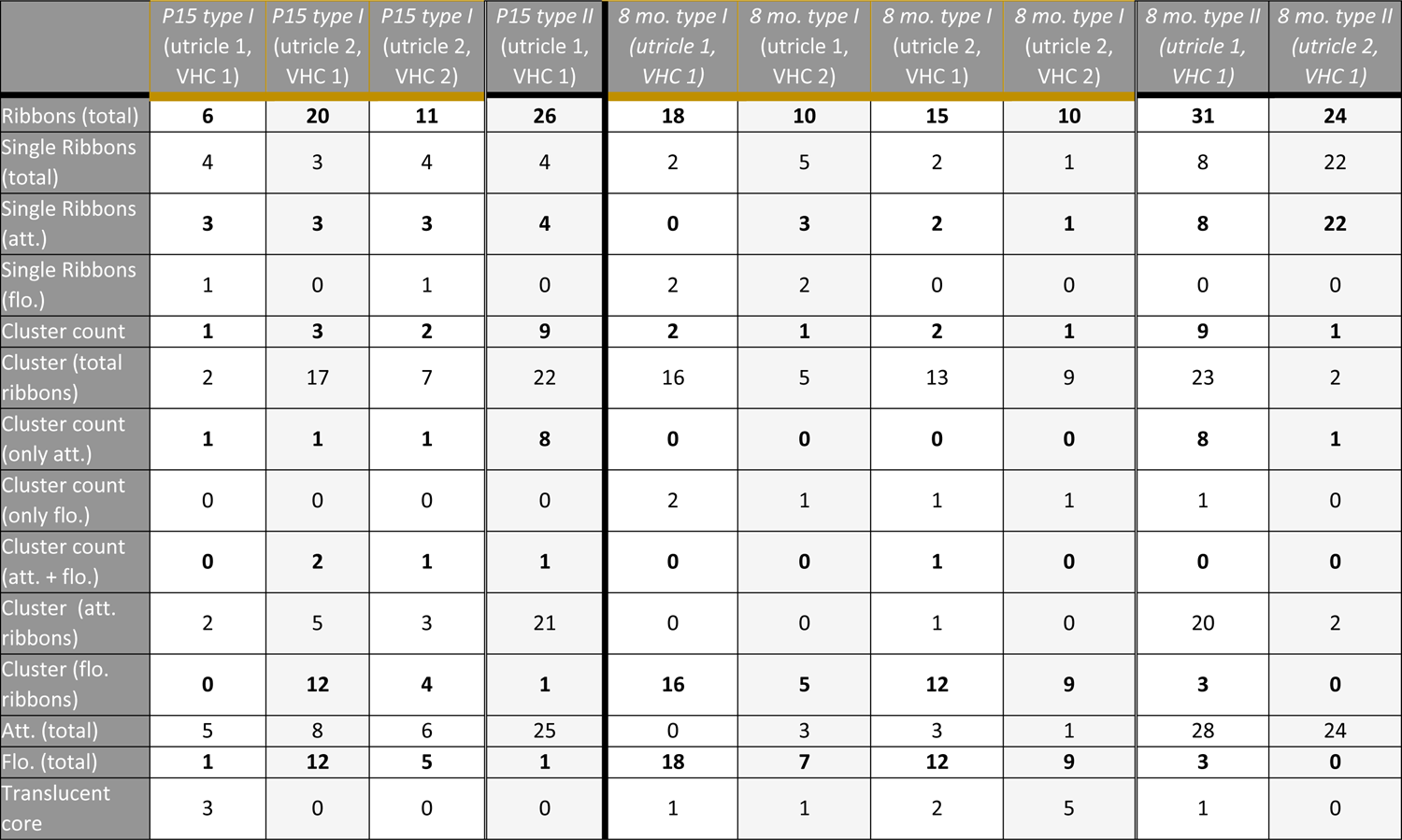
Counts of different ribbon categories from FIB-SEM. Absolute ribbon counts per VHC (Supplementary data for Fig. 4). Att. = attached, flo. = floating. P15 type I = 3 VHCs, N = 2 animals; P15 type II = 1 VHC, N = 1 animal; 8 mo. type I = 4 VHCs, N = 2 animals; 8 mo. type II = 2 VHCs, N = 2 animals.

**Table S6:**
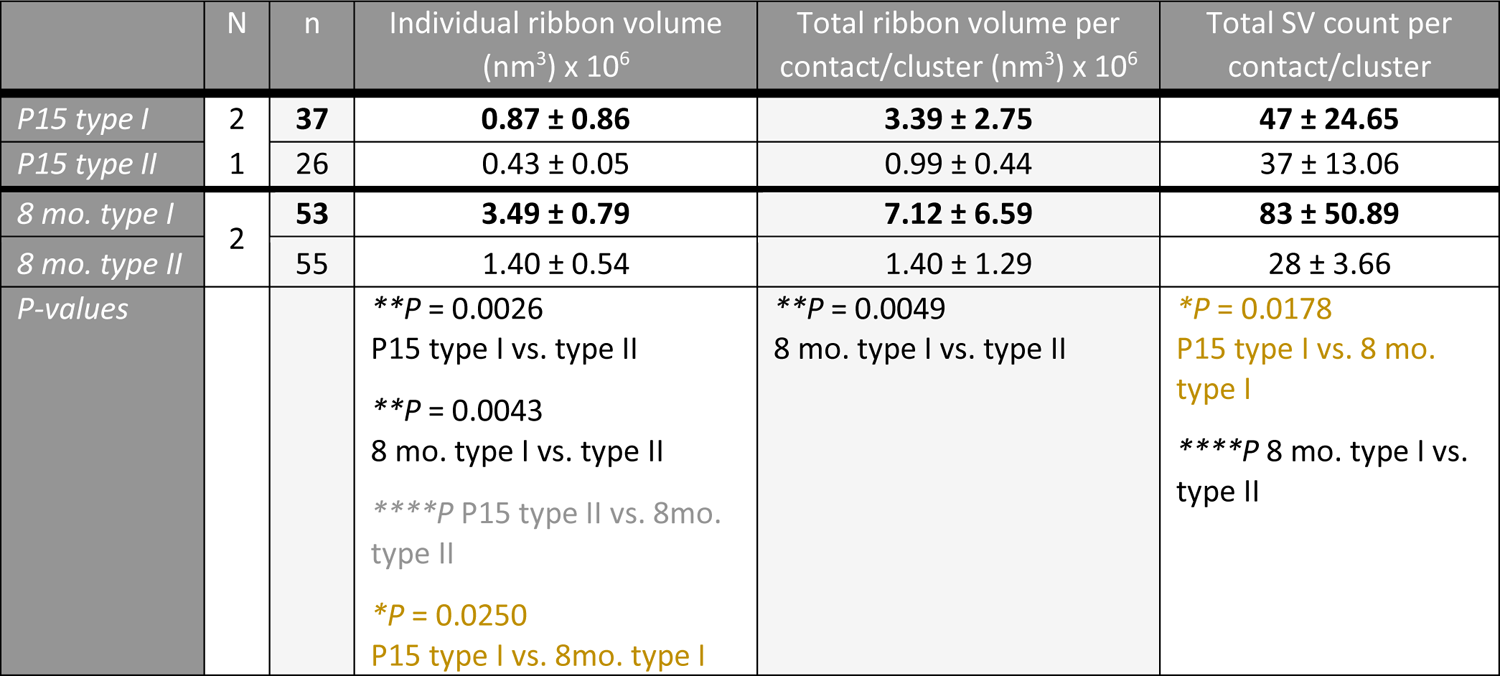
FIB-SEM data on SV count and ribbon size measurements. Ribbon size values and SV counts from type I and type II VHCs, respectively (Supplementary data for Fig. 4). Data are presented as median ± SEM. Significant results are highlighted with *\*P* < 0.05, *\*\*P* < 0.01, *\*\*\*P* < 0.001, *\*\*\*\*P* < 0.0001. N = animal count, n = ribbon count.

|  | N | n | Individual ribbon volume<br>(nm<sup>3</sup>) x 10<sup>6</sup> | Total ribbon volume per<br>contact/cluster (nm<sup>3</sup>) x 10<sup>6</sup> | Total SV count per<br>contact/cluster |
| --- | --- | --- | --- | --- | --- |
| P15 type I | 2 | 37 | 0.87 ± 0.86 | 3.39 ± 2.75 | 47 ± 24.65 |
| P15 type II | 1 | 26 | 0.43 ± 0.05 | 0.99 ± 0.44 | 37 ± 13.06 |
| 8 mo. type I | 2 | 53 | 3.49 ± 0.79 | 7.12 ± 6.59 | 83 ± 50.89 |
| 8 mo. type II |  | 55 | 1.40 ± 0.54 | 1.40 ± 1.29 | 28 ± 3.66 |
| P-values |  |  | **\*\*P = 0.0026**<br>P15 type I vs. type II<br>**\*\*P = 0.0043**<br>8 mo. type I vs. type II<br>**\*\*\*\*P** P15 type II vs. 8mo.<br>type II<br>**\*P = 0.0250**<br>P15 type I vs. 8mo. type I | **\*\*P = 0.0049**<br>8 mo. type I vs. type II | **\*P = 0.0178**<br>P15 type I vs. 8 mo.<br>type I<br>**\*\*\*\*P** 8 mo. type I vs.<br>type II |

**Table S7:** TEM data on SV counts. Number of SVs per ribbon from type I and type II VHCs, respectively (Supplementary data for Fig. 5 and Fig. S4). Data are presented as median ± SEM. Depending on the normality and the equality of variances tests, p-values are calculated by the two-tailed unpaired Student’s t-test or the unpaired two-tailed Mann-Whitney-Wilcoxon test between two samples. For multiple comparisons, the one-way ANOVA with post-hoc Tukey’s test or the KW test with multiple comparison correction was utilized. Non-significant differences are highlighted with *n.s.* and significant results are highlighted with *\*P* < 0.05, *\*\*P* < 0.01, *\*\*\*P* < 0.001, *\*\*\*\*P* < 0.0001. N = animal count, n = ribbon count.

|  | N | n | SV count per ribbon |  |  |  | SV density per ribbon surface (x 10<sup>-3</sup>) Attached | SV density per ribbon surface (x 10<sup>-3</sup>) Floating | Fraction per ribbon |  |
| --- | --- | --- | --- | --- | --- | --- | --- | --- | --- | --- |
|  |  |  | Attached | Floating | MP-SV | RA-SV |  |  | MP-SV | RA-SV |
| P9 type I | 2 | 86 | 13.0 ± 0.74 | 13.0 ± 0.67 | 3.0 ± 0.20 | 11.0 ± 0.70 | 0.66 ± 0.10 | 0.89 ± 0.10 | 0.22 ± 0.02 | 0.78 ± 0.02 |
| P9 type II |  | 63 | 9.0 ± 0.64 | 11.5 ± 0.98 | 3.0 ± 0.18 | 7.0 ± 0.57 | 0.69 ± 0.05 | 0.82 ± 0.14 | 0.28 ± 0.02 | 0.72 ± 0.02 |
| P15 type I | 2 | 128 | 9.0 ± 0.65 | 12.0 ± 1.53 | 2.0 ± 0.13 | 7.0 ± 0.57 | 0.68 ± 0.10 | 0.76 ± 0.10 | 0.22 ± 0.02 | 0.78 ± 0.02 |
| P15 type II |  | 86 | 9.0 ± 0.63 | 12.0 ± 0.44 | 2.0 ± 0.13 | 6.0 ± 0.53 | 0.73 ± 0.05 | 0.73 ± 0.10 | 0.27 ± 0.01 | 0.73 ± 0.01 |
| P20 type I | 2 | 165 | 14.0 ± 1.23 | 15.0 ± 0.90 | 3.0 ± 0.23 | 10.0 ± 1.30 | 1.08 ± 0.13 | 0.96 ± 0.00 | 0.18 ± 0.03 | 0.82 ± 0.03 |
| P20 type II |  | 72 | 9.0 ± 0.53 | 11.0 ± 0.72 | 2.0 ± 0.16 | 7.0 ± 0.46 | 0.75 ± 0.00 | 0.74 ± 0.18 | 0.24 ± 0.02 | 0.76 ± 0.02 |
| 3 mo. type I | 2 | 219 | 13.0 ± 1.09 | 16.0 ± 0.76 | 3.0 ± 0.30 | 9.0 ± 1.00 | 0.60 ± 0.10 | 0.56 ± 0.09 | 0.25 ± 0.02 | 0.75 ± 0.02 |
| 3 mo. type II |  | 151 | 10.0 ± 0.27 | 11.0 ± 0.60 | 3.0 ± 0.09 | 7.0 ± 0.26 | 0.75 ± 0.04 | 0.88 ± 0.08 | 0.27 ± 0.01 | 0.73 ± 0.01 |
| 9-11 mo. type I | 2 | 169 | 13.0 ± 1.31 | 14.0 ± 1.58 | 4.0 ± 0.50 | 10.0 ± 1.11 | 0.69 ± 0.17 | 0.92 ± 0.15 | 0.26 ± 0.04 | 0.74 ± 0.04 |
| 9-11 mo. type II |  | 53 | 10.0 ± 0.49 | 9.0 ± 1.13 | 3.0 ± 0.16 | 7.0 ± 0.38 | 0.66 ± 0.04 | 0.84 ± 0.10 | 0.29 ± 0.01 | 0.71 ± 0.01 |
| P-values |  |  | ***P = 0.0005<br>P9 type I vs. type II<br>**P = 0.0035<br>P20 type I vs. type II<br>***P = 0.0007<br>3 mo. type I vs. type II<br>*P = 0.0145<br>P15 type I vs. P20 type I<br>*P = 0.0331<br>P15 type I vs. 3 mo. type I<br>**P = 0.0091<br>P15 type I vs. P9 type I | ****P<br>3 mo. type I vs. type II<br>**P = 0.0017<br>P20 type I vs. type II<br>*P = 0.0459<br>P15 type I vs. 3 mo. type I | *P = 0.040<br>3 mo. type I vs. type II<br>*P = 0.0160<br>3 mo. type I vs. P15 type I | ***P = 0.0004<br>P9 type I vs. type II<br>**P = 0.002<br>3 mo. type I vs. type II<br>*P = 0.021<br>P20 type I vs. type II<br>*P = 0.0281<br>P9 type I vs. P15 type I<br>*P = 0.0214<br>P20 type I vs. P15 type I | n.s. | *P = 0.0486<br>P20 type I vs. 3 mo. type I<br>*P = 0.0213<br>3 mo. type I vs. type II | *P = 0.0193<br>P9 type I vs. type II | *P = 0.0193<br>P9 type I vs. type II |

**Table S8:**
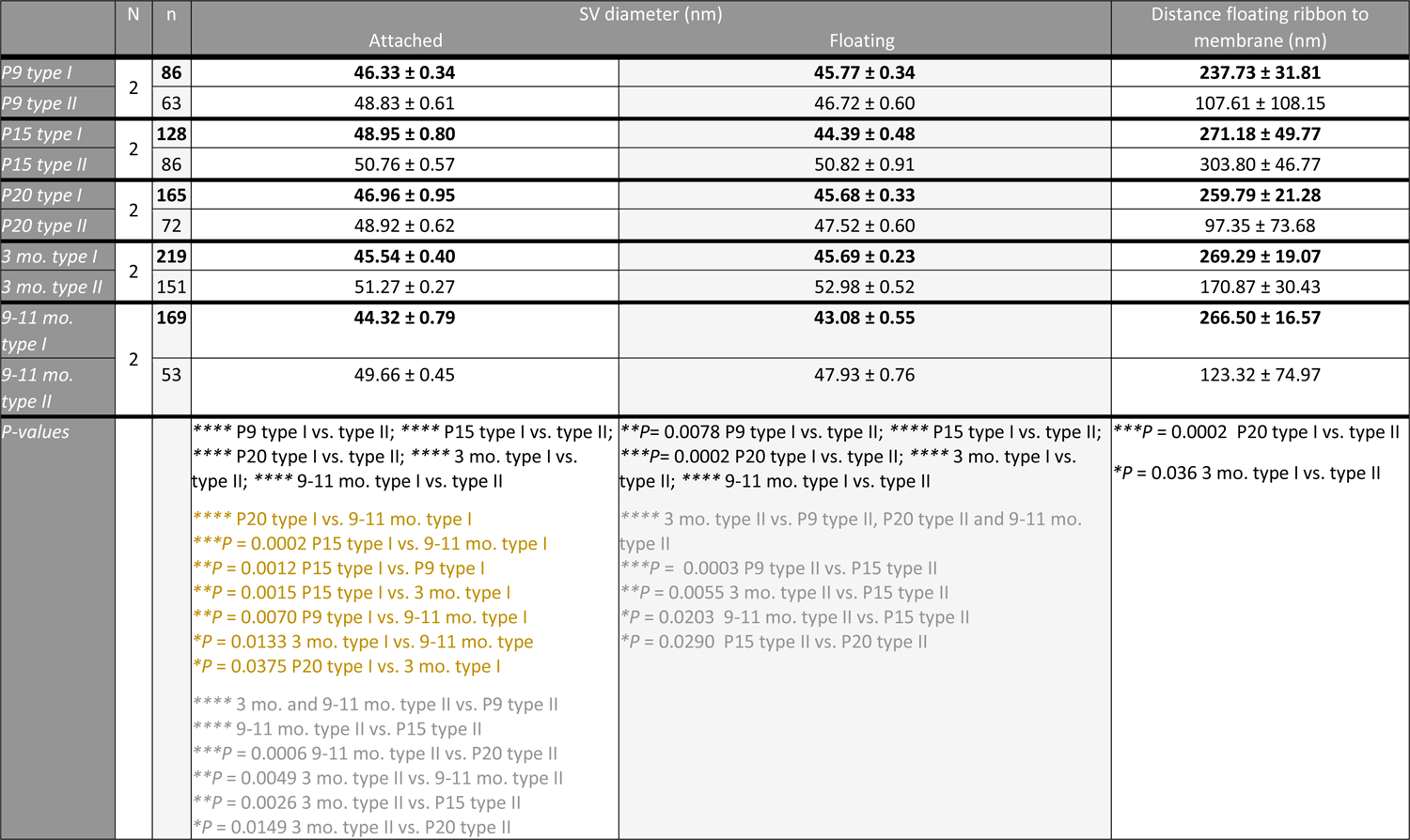
Size and distance measurements of AZ parameters. SV size and floating ribbon distance values at type I and type II VHCs (Supplementary data for Fig. 5 and Fig. S1). Data are presented as median ± SEM. Significant results are highlighted with *\*P* < 0.05, *\*\*P* < 0.01, *\*\*\*P* < 0.001, *\*\*\*\*P* < 0.0001. N = animal count, n = ribbon count. P-values are calculated by the two-tailed unpaired Student’s t-test or the unpaired two-tailed Mann-Whitney-Wilcoxon test between two samples. For multiple group comparisons, the KW test with multiple test correction was utilized.

|  | N | n | SV diameter (nm) |  | Distance floating ribbon to membrane (nm) |
| --- | --- | --- | --- | --- | --- |
|  |  |  | Attached | Floating |  |
| P9 type I | 2 | 86 | **46.33 ± 0.34** | **45.77 ± 0.34** | **237.73 ± 31.81** |
| P9 type II |  | 63 | 48.83 ± 0.61 | 46.72 ± 0.60 | 107.61 ± 108.15 |
| P15 type I | 2 | 128 | **48.95 ± 0.80** | **44.39 ± 0.48** | **271.18 ± 49.77** |
| P15 type II |  | 86 | 50.76 ± 0.57 | 50.82 ± 0.91 | 303.80 ± 46.77 |
| P20 type I | 2 | 165 | **46.96 ± 0.95** | **45.68 ± 0.33** | **259.79 ± 21.28** |
| P20 type II |  | 72 | 48.92 ± 0.62 | 47.52 ± 0.60 | 97.35 ± 73.68 |
| 3 mo. type I | 2 | 219 | **45.54 ± 0.40** | **45.69 ± 0.23** | **269.29 ± 19.07** |
| 3 mo. type II |  | 151 | 51.27 ± 0.27 | 52.98 ± 0.52 | 170.87 ± 30.43 |
| 9-11 mo. type I | 2 | 169 | **44.32 ± 0.79** | **43.08 ± 0.55** | **266.50 ± 16.57** |
| 9-11 mo. type II |  | 53 | 49.66 ± 0.45 | 47.93 ± 0.76 | 123.32 ± 74.97 |
| P-values |  |  | **** P9 type I vs. type II; **** P15 type I vs. type II; **** P20 type I vs. type II; **** 3 mo. type I vs. type II; **** 9-11 mo. type I vs. type II<br>**** P20 type I vs. 9-11 mo. type I<br>***P = 0.0002 P15 type I vs. 9-11 mo. type I<br>**P = 0.0012 P15 type I vs. P9 type I<br>**P = 0.0015 P15 type I vs. 3 mo. type I<br>**P = 0.0070 P9 type I vs. 9-11 mo. type I<br>*P = 0.0133 3 mo. type I vs. 9-11 mo. type I<br>*P = 0.0375 P20 type I vs. 3 mo. type I<br>**** 3 mo. and 9-11 mo. type II vs. P9 type II<br>**** 9-11 mo. type II vs. P15 type II<br>***P = 0.0006 9-11 mo. type II vs. P20 type II<br>**P = 0.0049 3 mo. type II vs. 9-11 mo. type II<br>**P = 0.0026 3 mo. type II vs. P15 type II<br>*P = 0.0149 3 mo. type II vs. P20 type II | **P= 0.0078 P9 type I vs. type II; **** P15 type I vs. type II; ***P= 0.0002 P20 type I vs. type II; **** 3 mo. type I vs. type II; **** 9-11 mo. type I vs. type II<br>**** 3 mo. type II vs. P9 type II, P20 type II and 9-11 mo. type II<br>***P = 0.0003 P9 type II vs. P15 type II<br>**P = 0.0055 3 mo. type II vs. P15 type II<br>*P = 0.0203 9-11 mo. type II vs. P15 type II<br>*P = 0.0290 P15 type II vs. P20 type II | ***P = 0.0002 P20 type I vs. type II<br>*P = 0.036 3 mo. type I vs. type II |

